# Lineage-Specific Immune Remodeling Following Autologous Hematopoietic Stem Cell Transplantation in Relapsing–Remitting Multiple Sclerosis

**DOI:** 10.1101/2025.07.01.662494

**Authors:** Sonia Gavasso, Jonas Bull Haugsøen, Dimitrios Kleftogiannis, Yola Gerking, Shamundeeswari Anandan, Ida Herdlevær, Håkon Olsen, Morten Brun, Nello Blaser, Bjørn Tore Gjertsen, Astrid Marta Olsnes, Einar Klæboe Kristoffersen, Kjell-Morten Myhr, Øivind Torkildsen, Lars Bø, Anne Kristine Lehmann

**Affiliations:** Department of Clinical Medicine, University of Bergen, Bergen, Norway; Neuro-SysMed, Department of Neurology, Haukeland University Hospital, Bergen, Norway; Department of Mathematics, University of Bergen, Bergen, Norway; Department of Informatics, University of Bergen, Bergen, Norway; Department of Clinical Science, University of Bergen, Bergen, Norway; Department of Medicine, Section of Hematology, Haukeland University Hospital, Bergen, Norway; Department of Immunology and Transfusion Medicine, Haukeland University Hospital, Bergen, Norway

## Abstract

**Background and Objectives:** Autologous hematopoietic stem cell transplantation (aHSCT) induces durable remission in treatment-refractory relapsing–remitting multiple sclerosis (RRMS). However, to what extent post-transplant immune recovery reflects restoration of pre-treatment state versus selective remodeling remains incompletely understood. We used high-dimensional whole-blood immune profiling to characterize lineage-specific remodeling in patients with RRMS following aHSCT and to compare trajectories with healthy reference profiles.

**Methods:** Peripheral blood from 25 patients with RRMS treated with aHSCT in the RAM-MS trial was analyzed longitudinally at baseline, 100 days, and 6, 12, and 24 months post-transplant. Samples were compared with those from 9 age- and sex-matched healthy controls using a 41-marker mass cytometry panel. Unsupervised clustering and multi-level modeling were used to quantify lineage-level abundance changes and phenotypic remodeling across adaptive and innate compartments.

**Results:** Immune recovery after aHSCT was characterized by selective and sustained reshaping of specific T-and B-cell subsets rather than uniform restoration of pre-treatment composition. Naïve CD4+ and CD8+ T cells gradually repopulated, whereas memory CD4+ T cells remained reduced at 24 months. Memory CD8+ T cells showed a transient early expansion before declining toward baseline levels. Naïve B cells expanded early and remained elevated, while memory B cells were durably reduced. B-cell reconstitution involved expansion of activation-associated naïve populations alongside contraction of selected memory subsets. Across T-cell compartments, CD161-expressing chemokine receptor–enriched subsets were selectively depleted. Several of these alterations persisted despite partial normalization of overall cell counts. These trajectories were consistent across patients, supporting a robust and reproducible effect of aHSCT on immune architecture.

**Discussion:** aHSCT induces durable, lineage-specific reorganization of adaptive immunity along a shared trajectory across patients, characterized by contraction of inflammatory T-cell programs and hierarchical restructuring of B-cell maturation rather than simple immune restoration.

## Introduction

Multiple sclerosis (MS) is a chronic inflammatory and degenerative disease of the central nervous system characterized by immune-mediated tissue injury and progressive neurological disability (1,2). Autologous hematopoietic stem cell transplantation (aHSCT) is among the most effective treatment options for patients with highly active relapsing-remitting MS (RRMS) who experience disease activity despite conventional disease-modifying therapies (3–6). Clinical trials and long-term observational studies have demonstrated sustained remission and durable suppression of inflammatory disease activity in a substantial proportion of aHSCT treated patients (7–10). However, it remains unclear whether long-term remission reflects quantitative recovery of immune cell counts or durable reorganization of adaptive immune architecture.

aHSCT induces profound lymphodepletion followed by gradual immune reconstitution. Previous studies using flow cytometry and repertoire sequencing have demonstrated contraction in memory T and B cell compartments, expansion of naïve populations, and renewal of the T cell receptor repertoire (11–17). However, prior work has largely focused on selected subsets or limited marker panels, and longitudinal high-dimensional profiling of whole blood immune composition remains limited.

High-dimensional mass cytometry enables quantification of multiple phenotypes simultaneously at single-cell resolution. Applied longitudinally, this approach allows comprehensive tracking of immune reconstitution across adaptive and innate compartments and may identify cellular subsets that persistently deviate from healthy reference profiles.

In this study, we performed 41-marker mass cytometry profiling of peripheral whole blood samples from 25 patients with RRMS treated with aHSCT in the RAM-MS trial, with longitudinal assessment from baseline to 24 months post-transplant and comparison with healthy controls. We aimed to define lineage-specific remodeling of immune architecture during long-term recovery following aHSCT.

## Methods

### Study design and cohort characteristics

This study included 25 patients with RRMS who underwent aHSCT in the Randomized Autologous heMatopoietic Stem Cell Transplantation Versus Alemtuzumab, Cladribine or Ocrelizumab for RRMS (RAM-MS) clinical trial between 2018 and 2021 (NCT03477500; EudraCT 2014-510630-40-00). All patients fulfilled the 2017 revised McDonald criteria for RRMS. Patients were enrolled based on the evidence of inflammatory disease activity during the year prior to inclusion despite receiving disease-modifying therapies (natalizumab, dimethyl fumarate, teriflunomide, fingolimod, interferon beta, or glatiramer acetate). Disease activity was defined as a clinical relapse with at least one new T1 contrast-enhancing lesion or three or more new or enlarging T2 lesions. The treatment protocol consisted of stem cell mobilization with cyclophosphamide and granulocyte colony-stimulating factor (G-CSF), followed by stem cell harvesting, lymphoablative conditioning with cyclophosphamide and antithymocyte globulin (ATG), and subsequent autologous hematopoietic stem cell transplantation (HSCT) (eFigure 1).

Baseline cohort characteristics are summarized in Table 1. The mean patient age was 38 years (16 females and 9 males). None of the patients had received prior T- or B-cell–depleting therapies, and no clinical relapses occurred during the protocol-defined drug-specific washout period prior to baseline sampling (ClinicalTrials.gov: NCT03477500).

**Table 1.** Baseline demographic and clinical characteristics of the 25 patients with RRMS included in the study. Ages are presented in ranges to preserve participant anonymity. 16 patients were female and 9 male. Previous treatments, protocol-defined washout prior to inclusion and baseline immune profiling, Expanded Disability Status Scale (EDSS) scores, and time since last relapse are shown. *In addition to the washout period, patients on teriflunomide received a washout protocol of cholestyramine 8 g three times daily for 11 days.

| *Patient* | *Age range (y)* | *Years since diagnosis* | *EDSS at inclusion* | *Previous treatment* | *Days from discontinuation of previous DMT to reinfusion* | *Days from qualifying MRI to reinfusion* |
| --- | --- | --- | --- | --- | --- | --- |
| 1 | 30-39 | 6 | 1 | Dimethyl fumarate | 79 | 191 |
| 2 | 30-39 | 9 | 1 | Teriflunomide | 48* | 63 |
| 3 | 40-49 | 5 | 1.5 | Natalizumab | 168 | 177 |
| 4 | 20-29 | 2 | 2.5 | Fingolimod | 203 | 214 |
| 5 | 20-29 | 1 | 1 | Teriflunomide | 107* | 239 |
| 6 | 20-29 | 5 | 2 | Dimethyl fumarate | 121 | 131 |
| 7 | 30-39 | 6 | 2 | Teriflunomide | 168* | 182 |
| 8 | 40-49 | 7 | 6 | Fingolimod | 148 | 171 |
| 9 | 30-39 | 12 | 3 | Fingolimod | 102 | 193 |
| 10 | 30-39 | 1 | 3.5 | Fingolimod | 95 | 247 |
| 11 | 30-39 | 9 | 2 | Fingolimod | 142 | 142 |
| 12 | 40-49 | 17 | 3 | Interferon beta | 92 | 147 |
| 13 | 40-49 | 12 | 3.5 | Fingolimod | 133 | 189 |
| 14 | 30-39 | 5 | 3.5 | Fingolimod | 107 | 134 |
| 15 | 40-49 | 8 | 2 | Dimethyl fumarate | 70 | 83 |
| 16 | 40-49 | 2 | 4 | Glatiramer acetate | 64 | 111 |
| 17 | 30-39 | 3 | 1 | Teriflunomide | 70* | 156 |
| 18 | 40-49 | 2 | 2 | Teriflunomide | 113* | 114 |
| 19 | 50-59 | 8 | 1.5 | Teriflunomide | 51* | 132 |
| 20 | 30-39 | 2 | 2.5 | Dimethyl fumarate | 51 | 113 |
| 21 | 30-39 | 12 | 2 | Fingolimod | 176 | 179 |
| 22 | 30-39 | 2 | 1 | Dimethyl fumarate | 108 | 140 |
| 23 | 30-39 | 9 | 1.5 | Teriflunomide | 45* | 122 |
| 24 | 40-49 | 2 | 2 | Teriflunomide | 54* | 99 |
| 25 | 40-49 | 1 | 2.5 | Dimethyl fumarate | 66 | 66 |
| **Mean** | **38** | **6** | **2** |  | **103** | **149** |

Peripheral whole-blood samples obtained at baseline and after 100 days (d), 6 months (m), 12 m, and 24 m post-aHSCT were used for immune profiling. A sex- and age-matched cohort of nine healthy individuals (5 females, 4 males, mean age 38) served as controls (Fig. 1).

**Figure 1.**
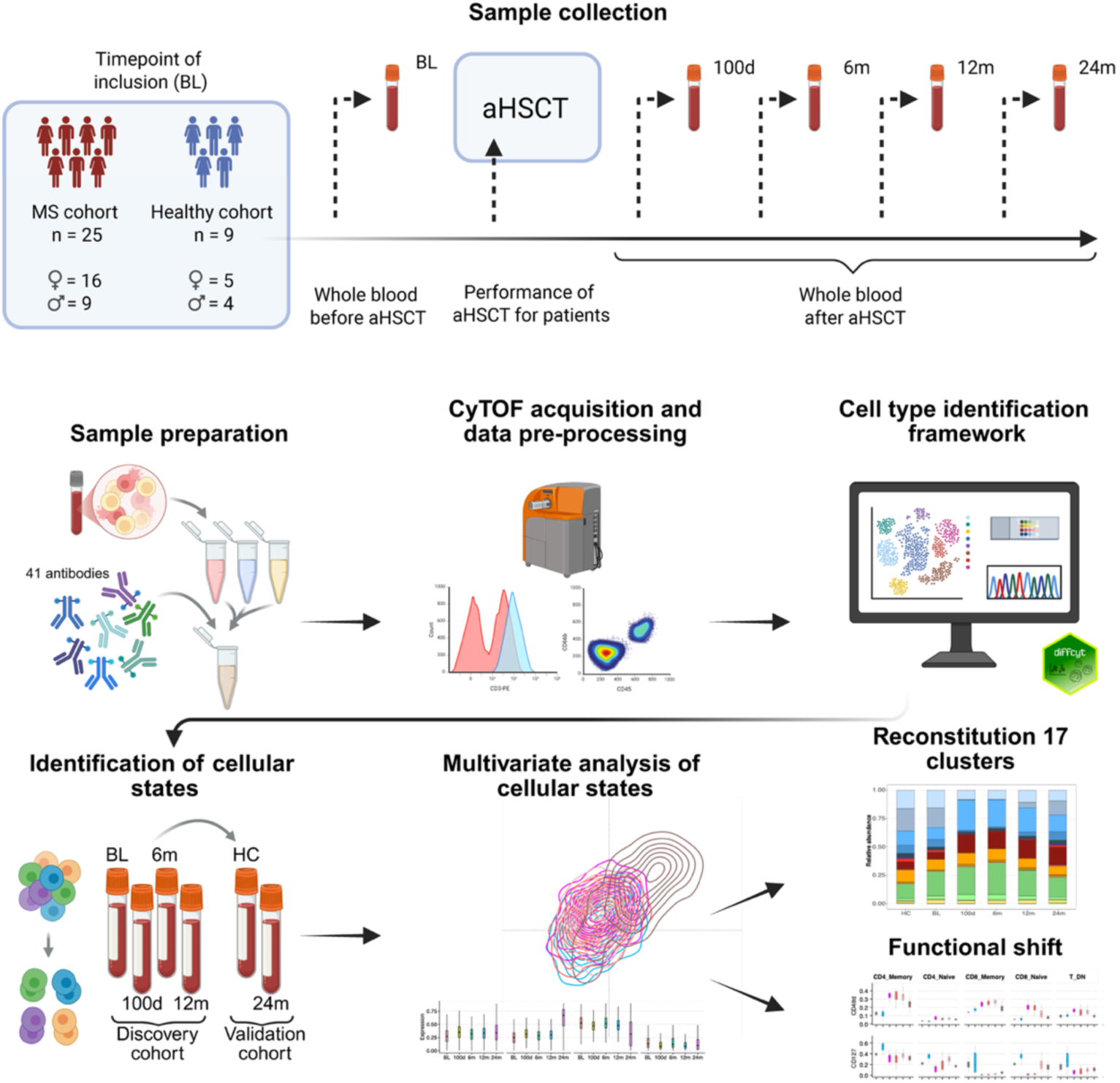
Sample collection, data acquisition and bioinformatics analysis. Sample collection. – Longitudinal sampling schedule for the MS cohort (n=25), consisting of patients who had recently experienced disease activity on highly effective disease-modifying therapies, and healthy controls (n=9). Whole blood samples were collected from all participants at baseline (BL). The MS patients underwent aHSCT, and additional samples were collected at 100d, 6m, 12m, and 24m post-transplant. **Sample preparation** consisted of barcoding of cells, sample pooling and staining with a 41-antibody panel. **Data acquisition** was performed using the CyTOF XT mass cytometer. **Data pre-processing** included cleanup, debarcoding, normalization, and initial gating on CD45 and CD66b for lymphocyte/granulocyte gating. For the **cell type identification framework**, 17 lymphocyte subpopulations were identified using FlowSOM clustering with metacluster merging via ConsensusClusterPlus, visualized with UMAP, and compared across conditions using differential abundance analysis with diffcyt to assess changes in cell population distribution before and after aHSCT. **Identification of cellular states** resulted in 107 cell states in the discovery cohort using FlowSOM clustering, ConsensusClusterPlus, and cell-type-specific LDA models, which were then used to predict these states in the temporal extension cohort (24 months and HC). Finally, **multivariate analysis** was performed for selected states using FreeViz visualizations and SARA analysis. Reconstitution dynamics were assessed by tracking temporal changes in relative abundances of the **17 identified lymphocyte clusters** from baseline (BL) to 24m post-aHSCT, accompanied by analysis of **functional shifts** illustrated by persistent changes in key marker expressions across lymphocyte populations, highlighting sustained immune reprogramming post-aHSCT.

### Standard Protocol Approvals, Registrations, and Patient Consents

The study was approved by the Regional Committee for Medical and Health Research Ethics, Western Norway (REK 606396), and all participants provided written informed consent.

### Mass cytometry profiling and data preprocessing

Peripheral whole-blood samples were stabilized within 15-20 minutes after collection to preserve the *in-vivo* cellular composition and minimize handling-induced artifacts. To ensure longitudinal comparability, samples from each patient were processed within shared barcode pools whenever possible. Cells were barcoded, pooled, and stained with a 41-parameter mass cytometry panel (eTable 1) specifically designed for the RAM-MS aHSCT study to capture lineage identity together with trafficking, activation, differentiation, and homeostatic markers relevant to immune reconstitution (18–20) (see eMethods for a detailed description of the mass cytometry and data preprocessing).

Antibody titration and panel optimization were performed prior to study initiation to minimize signal spillover and ensure robust detection across all marker ranges. Data were acquired on a CyTOF XT instrument using standardized procedures incorporating EQ calibration beads and a shared reference control sample within each barcode pool to enable intra- and inter-run signal normalization. Longitudinal samples were acquired under harmonized instrument settings to minimize acquisition drift across time points.

Following bead normalization and debarcoding, doublets and debris were excluded based on DNA intercalator intensity and event length. CD45^high^, CD66b^−^ leukocytes were retained for downstream analysis to focus on mononuclear immune compartments, while preserving granulocyte proportions for quality control (eFigure 2). Marker intensities were arcsinh-transformed prior to clustering. Residual inter-run variation was minimized by reference control–based signal alignment across acquisition batches.

To confirm that stabilization and fixation did not distort leukocyte distributions, CyTOF-derived CD45^high^, CD66b^−^ and granulocyte fractions were compared with routine clinical differential counts obtained from fresh blood samples. Cell retention numbers, batch metrics, additional quality control analyses, detailed acquisition parameters, antibody panel composition, normalization procedures, and preprocessing workflows are described in the eMethods.

### Bias and measures to minimize bias

Several potential sources of bias were considered. Selection bias was limited by including all available aHSCT-treated participants from the RAM-MS trial with sufficient biological material for longitudinal immune profiling; however, the sample size was constrained by material availability and should therefore be considered a limitation. Technical bias was minimized by rapid whole-blood stabilization, shared barcode pooling of longitudinal samples whenever possible, standardized CyTOF acquisition, EQ bead normalization, reference-control–based batch alignment, antibody titration, and harmonized instrument settings. Analytical bias was reduced by using unsupervised clustering for lineage identification, predefined preprocessing and quality-control criteria, paired statistical models accounting for repeated measures, false discovery rate correction, and no imputation of missing samples. To reduce bias from time-point–specific clustering, functional cellular states were defined in a discovery cohort and then projected onto later time points and healthy controls using trained lineage-specific models. Interpretation bias was further limited by applying orthogonal analytical approaches, including FreeViz and SARA, to support findings derived from abundance and state-level analyses. Despite these measures, residual bias related to the small cohort size, incomplete sampling at some time points, and the absence of randomized healthy controls cannot be excluded.

### Data analysis

To characterize immune reconstitution beyond simple numeric recovery, we applied a multi-layer analytical framework designed to resolve remodeling at progressive levels of granularity. This approach comprised: (i) lineage-level abundance reconstitution, (ii) the definition and longitudinal tracking of persistent functional cellular states within those lineages, and (iii) a higher order assessment of the functional displacement axes at the single-cell level. By utilizing multivariate metrics (FreeViz and SARA) to support our primary findings, this framework enabled formal separation of quantitative repopulation from internal architectural remodeling and coordinated phenotypic reprogramming (see eMethods for a detailed description of the computational pipeline and multivariate metrics).

Unsupervised clustering using FlowSOM followed by consensus meta-clustering was performed within the CD45^high^, CD66b^−^ compartment to identify major immune populations (21,22). Seventeen populations were annotated based on the expression of canonical lineage markers and included CD4^+^ and CD8^+^ T-cell subsets, B-cell subsets, NK-cell subsets, and monocyte/dendritic cell subsets. Global variation in immune composition across time points was visualized by PCA using per-sample relative abundances of the 17 populations.

Differential abundance across time points was assessed using the diffcyt-DA-edgeR framework with paired modeling to account for repeated measurements within individuals (23). False discovery rate correction was applied, and log₂ fold changes were calculated relative to baseline to define lineage-level reconstitution kinetics. Missing samples were not imputed; paired models and per-time-point summary statistics included all available observations.

To assess remodeling within the repopulating lineages, multivariate functional cellular states were defined within each of the 17 immune populations. In a discovery cohort (baseline–12 m), unsupervised clustering based on 28 additional functional markers, including chemokine receptors, adhesion molecules, activation markers, and differentiation-associated proteins, identified 107 reproducible cellular states. These states represent stable multivariate phenotypes rather than dataset-specific clusters. To ensure longitudinal consistency and avoid re-clustering of all samples, including later-acquired time points, lineage-specific linear discriminant models were trained on the discovery cohort and subsequently used to project state definitions onto the remaining discovery samples, as well as the temporal extension dataset (24 month time point) and external reference (healthy controls). All samples were jointly re-clustered at the lineage level to define the 17 canonical immune populations. Persistently altered states were defined based on sustained log₂ fold change deviations across consecutive post-treatment time points, thereby identifying durable architectural remodeling.

To provide orthogonal support for the coordinated phenotypic reprogramming observed, we utilized supervised dimensionality reduction and distributional shift analysis. Specifically, single-cell profiles from baseline and 24 months were projected using FreeViz (24) to visualize the global multivariate separation and identify the markers contributing most to the transition between these states. This supervised projection served to illustrate a lineage-specific functional remodeling axis.

Finally, to validate these findings at higher resolution, marker-level distributional shifts within persistently altered cellular states were assessed using SARA (Significance Analysis of Response to Activation) (25). By comparing full single-cell intensity distributions rather than mean values, SARA provided a confirmatory, state-specific measure of directional shifts (induction vs. inhibition) at 24 months. These analyses supplemented the primary framework by quantifying the degree of coordinated functional remodeling without influencing the initial state definitions.

All analyses were performed in R. The complete analysis pipeline, including scripts required to reproduce the computational workflow, is publicly available at https://github.com/dkleftogi/immuneReconstitution_RRMS.

### Data Availability

Anonymized CyTOF datasets generated during this study are available from the corresponding author upon reasonable request for research purposes, subject to clinical trial data protection regulations.

## Results

### Patient cohort

The cohort consisted of 25 patients (16 female, 9 male) with a mean age of 38.0 years (range 22–50). At the time of inclusion, patients had a mean disease duration of 6.0 years (range 1–17) and a median EDSS score of 2.0 (range 1.0–6.0). All patients were transitioning from previous disease-modifying therapies (DMTs), most commonly fingolimod (n=8), teriflunomide (n=8), and dimethyl fumarate (n=6). The mean washout period from previous DMT discontinuation to reinfusion was 103 days, and the mean interval from the qualifying MRI to reinfusion was 149 days (Table 1).

### Broad whole-blood immune composition is preserved, whereas individual lineages show distinct post-aHSCT reconstitution trajectories

To evaluate the long-term cellular impact of treatment, we analyzed longitudinal whole-blood CyTOF profiles from 25 patients with RRMS treated with lymphoablative aHSCT (cyclophosphamide/ATG) in the RAM-MS trial and compared them to 9 age- and sex-matched healthy controls (HC). Samples were collected at BL and at 100d, 6m, 12m, and 24m post-aHSCT.

Across time points, the relative proportions of granulocytes and CD45^high^CD66b^−^ leukocytes in fixed whole blood were broadly stable and comparable to HC (Fig. 2a). The leukocyte fraction correlated strongly with routine flow cytometry differential counts from fresh blood (R² = 0.90; eFigure 3), supporting robust recovery of expected leukocyte fractions.

**Figure 2.**
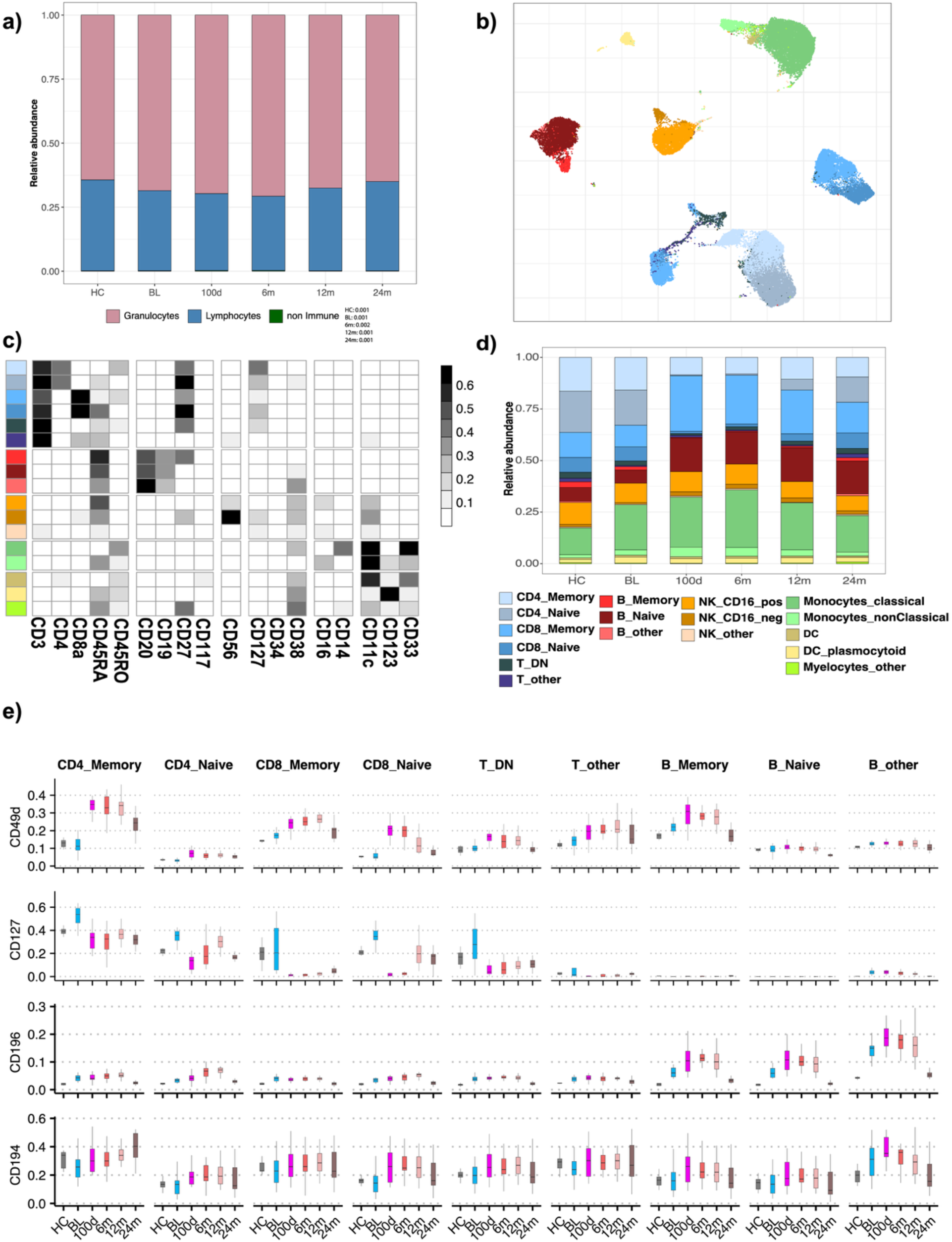
Immunophenotyping of cell populations. **a**, Relative abundance of manually gated populations at baseline (BL, pre-aHSCT), and at 100d, 6m, 12m, and 24m post-aHSCT, as well as in healthy controls (HC). Populations include CD45highCD66b-, CD45lowCD66b+, and CD45-CD66b- subsets. CD45lowCD66b+ cells were the predominant population (64.3–70.6%), followed by CD45highCD66b- cells (29–35.4%), with a minor fraction of CD45-CD66b- cells (0.15–0.27%). **b**, UMAP visualization of 17 clusters identified within the CD45highCD66b- gate using the FlowSOM / ConsensusClusterPlus framework. The plot was generated from 50,000 randomly selected cells (2,000 per patient) across 25 MS patients and time points. **c**, Heatmap showing average expression of 18 phenotypic markers used for clustering across the 17 identified cell populations. **d**, Stacked bar plots displaying the relative abundance of the 17 clusters within the CD45highCD66b- gate across all time points, with data from HCs shown for comparison. A consistent colour scheme is used in (b–d) to represent the same cell populations. **e**, Boxplots showing the expression of selected markers across identified T and B cell clusters. CD49d is an integrin involved in CNS migration and the target of anti-CD49d therapies in MS; CD127 is the IL-7 receptor involved in T cell survival and homeostasis; and CD196 (CCR6) directs CCR6+ Th17 cells to sites of CNS inflammation.

Within the CD45^high^CD66b^−^ leukocytes, unsupervised clustering identified 17 immune populations spanning T-, B-, NK-, monocyte, dendritic and residual myeloid compartments (Fig. 2b–d; eFigures 4–6). PCA of per-sample population abundances revealed a consistent shared shift at 100d, followed by gradual convergence toward HC profiles (eFigure 7). Major immune lineages and subpopulations were defined by manual annotation of FlowSOM meta-clusters based on median marker expression profiles. Five principal lineages and 17 subpopulations were identified, with marker definitions and cluster IDs provided in eTables 2A–B.

Within the 17 immune populations, reconstitution was highly structured and lineage-specific and was accompanied by sustained shifts in trafficking/homeostatic marker distributions. Beyond changes in abundance, several trafficking and homeostasis-associated markers showed durable deviations across the adaptive compartments (Fig. 2e; eFigure 8). CD49d increased broadly after aHSCT, peaking at 6–12m and remaining elevated at 24m in naïve and memory CD4 and CD8 T cells, while declining below HC levels in naïve B cells at 24m. Conversely, CD127 was reduced from BL onward across T-cell subsets and remained suppressed relative to HC at 24m. CCR6 was elevated at BL, increased early in naïve compartments post-aHSCT, and returned toward HC by 24m, whereas CCR4 increased post-aHSCT in selected T- and B-cell subsets, exceeding HC levels, particularly in CD4 memory cells.

### Coordinated reconstitution trajectories reveal persistent naïve T-cell lymphopenia and B-cell skewing

Across patients, population-level reconstitution followed highly consistent trajectories (Fig. 3a). Naïve CD4 and naïve CD8 T cells were markedly depleted early and recovered gradually; CD4 naïve T cells remained below BL and HC even at 24m, whereas CD8 naïve T cells approached BL/HC by 24m. CD4 memory T cells declined and remained persistently reduced at 24m. In contrast, CD8 memory T cells expanded at 100d and then gradually declined. T_DN decreased early but recovered to BL/HC levels by 24m, while T_other was comparatively stable and rose from below-HC BL levels toward HC levels by 24m.

**Figure 3.**
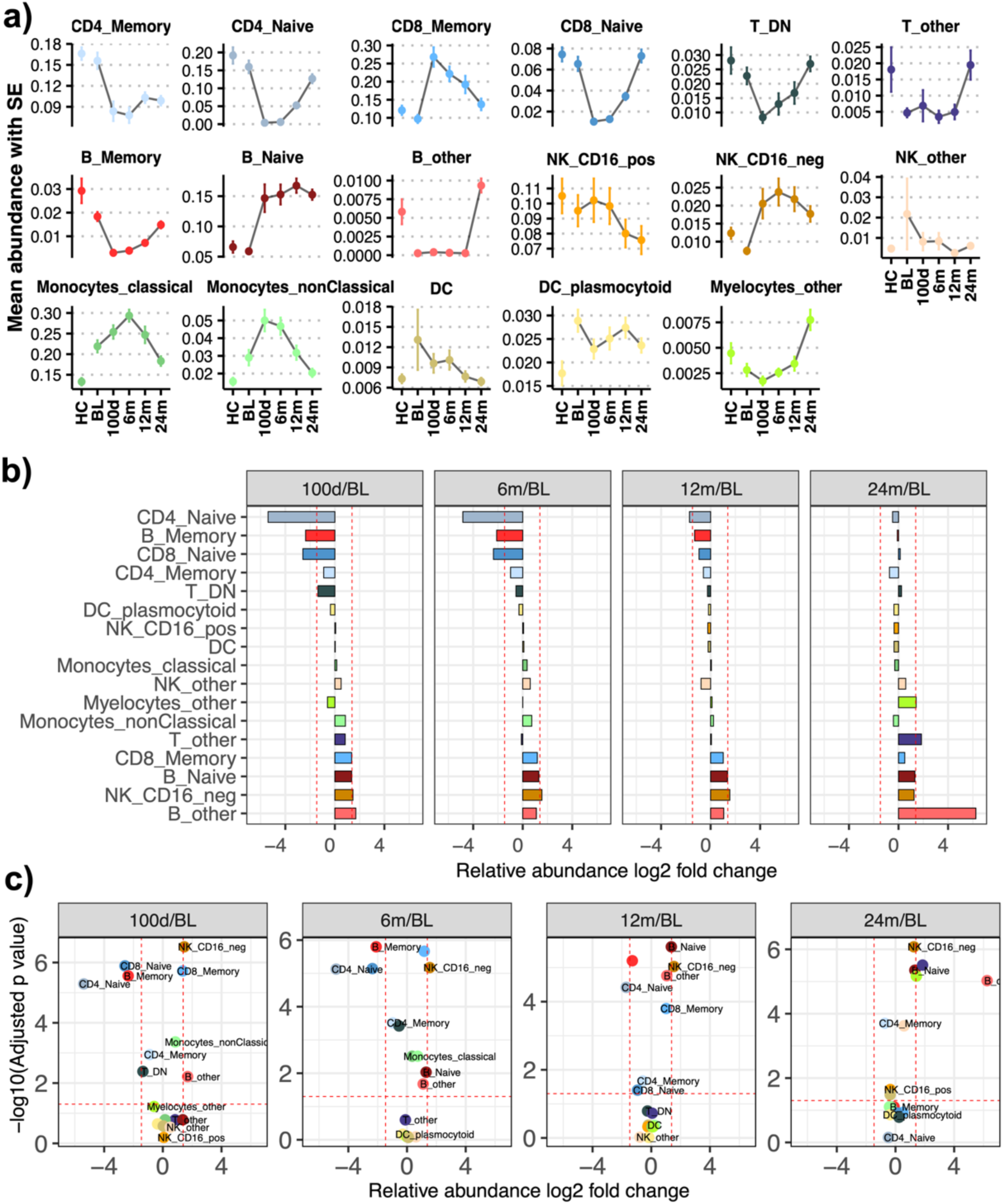
Immune reconstitution and differential abundance analysis pre- and post-aHSCT. **a**, Mean relative abundance of immune cell populations per patient ± standard error (SE). **b**, Model-estimated log₂ fold changes (log₂FC) from paired diffcyt-DA-edgeR analyses for each post-aHSCT time point relative to baseline (BL). Cell populations are ordered from highest to lowest average log_2_FC. **c**, Volcano plots summarizing differential abundance analysis using diffcyt-DA-edgeR. Statistical significance was defined as FDR-adjusted ***P < 0.05***. Vertical dashed lines indicate empirical log₂FC thresholds (−1.25 and +1.97), corresponding to the 10th and 90th percentiles of the pooled log₂FC distribution and shown for visualization of the most extreme abundance shifts only.

In the B-cell compartment, memory B cells were depleted after 100d and remained lower than HC at 24m, whereas naïve B cells expanded from 100d onward and remained elevated through 24m. NK CD16^−^ increased early and remained elevated at 24m, while NK CD16^+^ declined after 6m. Myeloid and DC populations showed smaller but persistent deviations, including sustained differences relative to HC at 24m (Fig. 3a).

Differential abundance modelling (diffcyt-DA-edgeR) confirmed significant depletion of naïve CD4, naïve CD8 T cells and memory B cells at 100d, persisting through 6–12m, with partial/variable recovery by 24m (Fig. 3b–c; eTable 3). Notably, several lineages remained significantly different from baseline and/or HC at 24m, indicating a transition away from the baseline state toward more HC-like profiles without full convergence (eFigure 9).

### Functional cellular states reveal durable remodeling within repopulating lineages

To resolve functional heterogeneity within the 17 immune populations, we defined 107 multivariate cellular states in a discovery cohort (BL–12m) using 28 additional markers and projected these states onto a temporal extension cohort (24m and HC) using supervised machine learning models (eFigures 10 and 11). While population abundances broadly converged toward HC (Fig. 4; eFigure 12), state-level analysis identified 17 persistently altered states (≥6m beyond empirical effect-size thresholds), concentrated within naïve lymphocyte compartments but also present in memory and innate-like T-cell populations (Fig. 5; eFigure 12).

**Figure 4.**
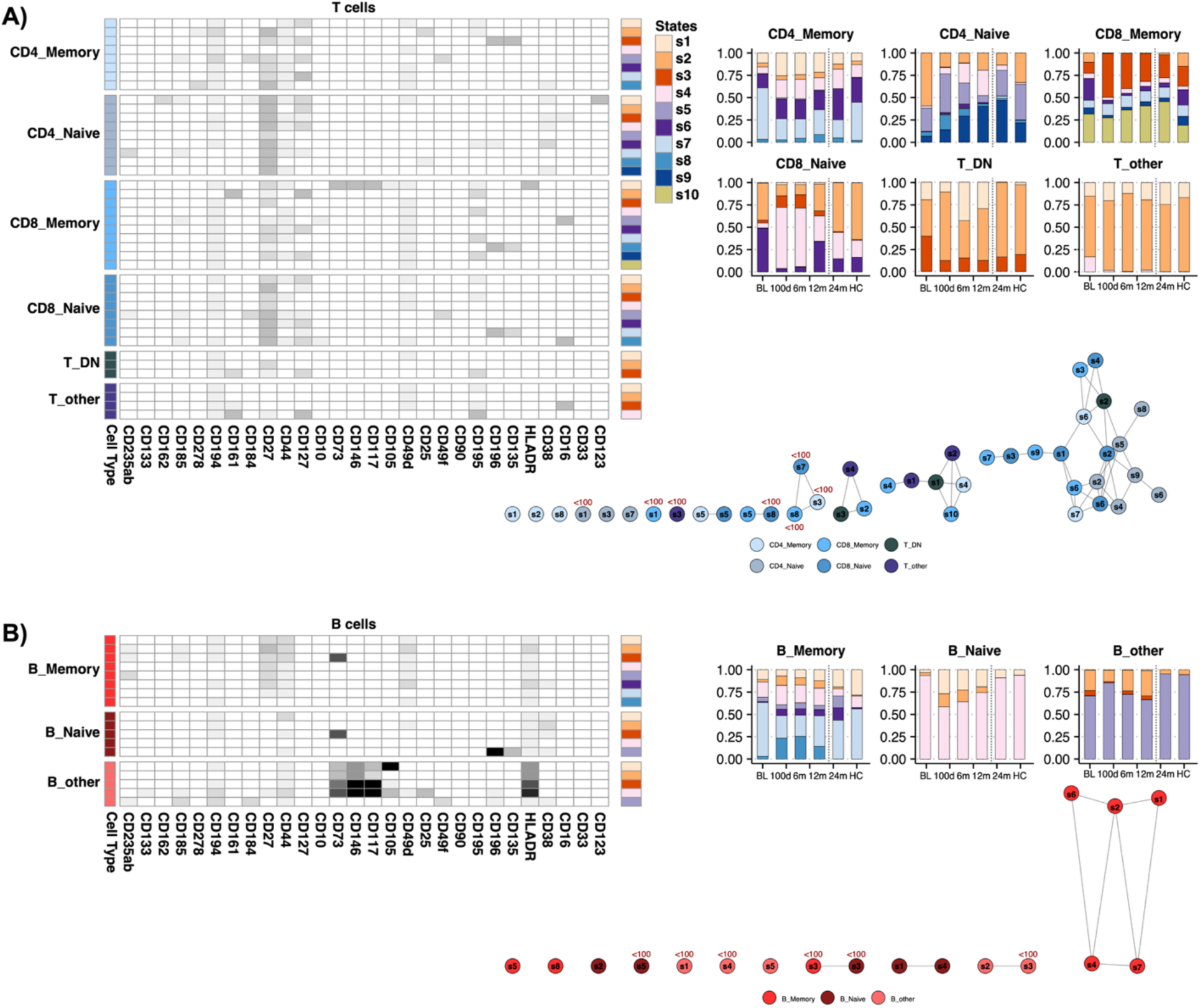
Functional cellular states identified and longitudinally mapped across adaptive immune populations. Left: Heatmap showing average expression profiles of T cell (**a.**) and B cell (**b.**) cellular states defined by unsupervised clustering within the discovery cohort (BL-12m), based on 28 additional markers. Right: Relative abundance of these predefined cellular states across all timepoints, including projected states in the temporal extension dataset (24m and HCs), assigned using the trained LDA models. States are displayed within their respective lineage-defined immune populations. This unified representation allows direct biological comparison of functional state dynamics before and after aHSCT, and relative to healthy controls. **Functional graphs of cellular state associations within each adaptive cellular compartment.** Graphs showing associations between the expression profiles of cellular states within each cellular compartment. Cellular states are connected by lines if their Pearson correlation exceeds 0.9. States containing fewer than 100 cells are specifically marked.

### Naïve T-cell compartments are rebuilt with altered internal composition

Consistent with lymphoablative conditioning, naïve CD4 and naïve CD8 T cell populations showed profound early depletion with gradual recovery (Fig. 3; Fig. 2d). However, their internal state composition did not simply revert to BL (Fig. 4 and 5).

**Figure 5.**
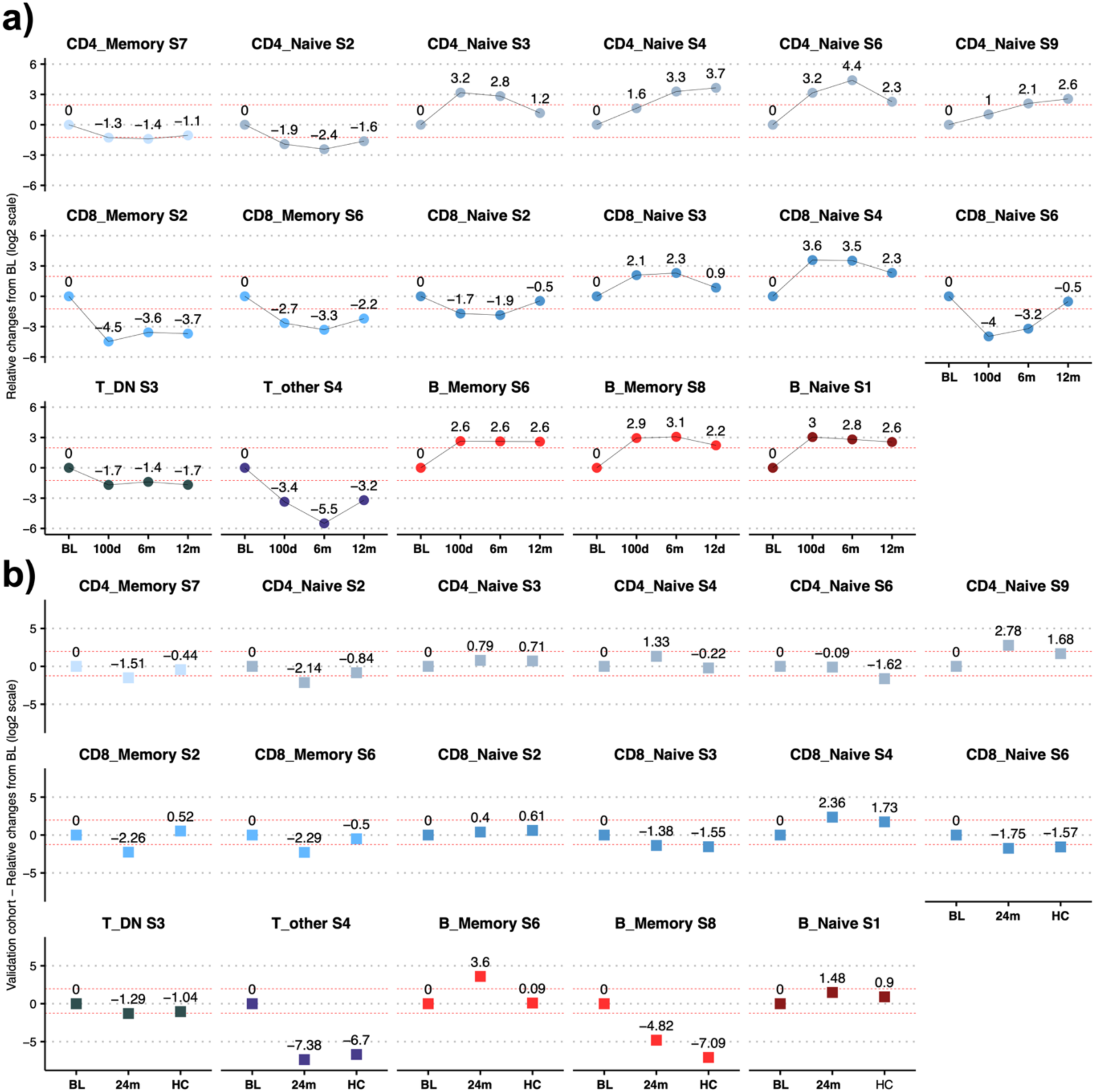
Persistent cellular state alterations following aHSCT. **a)** Discovery cohort (BL–12m): Dot plots display model-estimated log₂ fold changes (log₂FC) in the relative abundance of predefined cellular states across post-treatment time points relative to baseline. States shown are those exceeding the empirical 10th or 90th percentile thresholds (−1.25 and +1.97) and remaining beyond these thresholds for at least two consecutive post-treatment time points (≥6 months). Each dot represents the cohort-level log₂FC estimate for a given time point. Red horizontal lines indicate the percentile-derived effect size thresholds. These thresholds were used for biological interpretation and not for statistical inference. **b)** Temporal extension cohort (24m and HCs): The same predefined cellular states identified as persistently altered in the discovery cohort were projected onto the temporal extension dataset (24m and HCs) using trained LDA models. Dot plots show corresponding model-estimated log₂FC values relative to baseline. Red horizontal lines indicate the discovery-derived thresholds, applied without recalibration in the temporal extension cohort (24m and HC).

Within CD4 naïve T cell subset, one state (S2) remained persistently depleted at 24m (log₂FC −2.14) and was characterized by CD127^+^CD44^−^ expression with minimal expression of activation or trafficking markers, consistent with a quiescent IL-7Rα–dependent phenotype. In contrast, state S9 remained durably expanded (log₂FC +2.78), lacked the expression of CD127, but was enriched with CD44^+^ and CD38^+^, suggesting a more activated, turnover-associated naïve-like phenotype. A chemokine receptor–rich state (S6; CD127^−^, CD44^+^, CD38^+^ with CCR4/CCR5/CCR6 expression) expanded early but contracted over time, indicating selective long-term suppression of trafficking-enriched naïve phenotypes. In addition, a distinct CD25^+^ state (S8) expanded prominently at 100d but did not meet persistence criteria, suggesting a transient early enrichment of regulatory-associated naïve phenotypes during lymphopenic reconstitution.

Within CD8 naïve T cell subsets, one state expressing CD49d^+^, CD127^−^, CCR4^+^, CD44^−^ (S4) remained persistently elevated at 24m, while the reciprocal CD127^+^, CD49d^−^, CCR4^−^, CD44^+^state (S6) remained persistently reduced, indicating durable rebalancing away from homeostatic CD127-high naïve phenotypes toward CD49d-associated states.

Taken together, these findings demonstrate that aHSCT durably reshapes the internal architecture of the naïve T-cell compartment. Naïve T-cell recovery was accompanied by persistent shifts in state composition, including expansion of CD127-negative CD44/CD38- or CD49d-expressing subsets and reduction of CD127-high populations. Thus, reconstitution of the naïve compartment proceeded via sustained compositional remodeling rather than restoration of baseline state distributions.

### Selective depletion of CD127- and CD161-enriched programs within memory and innate-like T-cell compartments

At the population level, CD4 memory T cells remained persistently reduced at 24m, whereas CD8 memory T cells showed transient early expansion followed by gradual decline (Fig. 3a). At the state level, in CD4 memory T cells, one state met the persistence criteria: S7 (CD27^+^, CD44^+^, CD127^high^, CCR4^+^ and CD49d^+^) showed sustained depletion following aHSCT, consistent with the prolonged reduction in total CD4 memory T cell abundance at 24m and indicating selective contraction of a CD127^high^ trafficking-associated memory phenotype.

In CD8 memory T cells, two states were persistently reduced. S2, defined by CD161^high^, CD44^−^, CD27^low^, CD127^high^, CD49d^−^, CCR5^+^, declined durably, as did S6 (CD161^−^, CD44^+^, CD27^+^, CD127^+^, CD49d^+^, CCR4^+^). Thus, despite transient early expansion of total CD8 memory cells, state-level analysis revealed sustained depletion of distinct CD161^+^ and CD127^+^ memory programs.

Similarly, within the T_DN and T_other compartments, one state in each (T_DN S3 and T_other S4) declined markedly and persistently. Both were enriched for CD161^+^ and CD127^+^ and expressed CCR4^+^ and CCR5^+^, consistent with durable loss of chemokine receptor–rich innate-like T-cell states.

### Coordinated depletion of a CD161^high^ T-cell module across memory and innate-like compartments

Within the T-cell compartment, persistently depleted states formed a discrete structural cluster in the state network characterized by high CD161^+^ expression (Fig. 4 functional graphs). This triad comprised CD8_Memory S2 (CD161^high^, CCR4^−^, CD27^low^, CD127^high^, CCR5^+^), T_DN S3 (CD161 ^high^, CCR4^+^, CD27^low^, CD127^high^, CCR5^+^), and T_other S4 (CD161 ^high^, CCR4^+^, CD27^low^, CD127^high^, CCR5^+^). Each of these states showed sustained contraction following aHSCT. Importantly, these state-level contractions occurred within lineages that exhibited divergent population-level kinetics, underscoring that internal architectural remodeling was not simply a reflection of global lineage contraction. The convergence of these phenotypically related states across distinct T-cell compartments indicates selective depletion of a shared CD161^high^ chemokine receptor–rich T-cell module rather than compartment-restricted remodeling.

### B-cell compartments undergo reciprocal remodeling of naïve and memory states

In contrast to T-cell compartments, B-cell reconstitution was characterized by expansion of naïve cells and selective restructuring within memory subsets. At the population level, naïve B cells increased from 100d onward and remained elevated at 24m, whereas memory B cells were markedly depleted early after aHSCT and partially recovered over time. The B_other compartment remained persistently reduced relative to baseline and healthy controls (Fig. 3a).

Within the naïve B-cell compartment, three phenotypically distinct states were identified, but only one state (S1) met persistence criteria. S1 expanded early post-transplant and remained durably elevated at 24m (log₂FC +1.48 vs BL), approaching healthy control levels (Fig. 4 and 5). Phenotypically, S1 expressed CD44^+^, CCR4^+^ (CD194), HLA-DR^+^ and CD38^+^, consistent with an activation- and antigen-presentation–associated naïve B phenotype. Other naïve B states fluctuated but did not demonstrate sustained displacement.

In the memory B-cell compartment, two states exhibited persistent and reciprocal remodeling. S6 (CD27^+^, CD44^low^, CD194^+^, CD49d^high^, HLA-DR^+^, CD38^+^) expanded and remained elevated beyond 6m. In contrast, S8 (CD27^−^, CD44^−^, CD194^+^, CD49d^+^, HLA-DR^+^, CD38^−^) declined durably following aHSCT. Memory B-cell reconstitution was therefore characterized by differential representation of CD27^+^and CD27^−^ subsets rather than uniform recovery across memory populations. The sustained reduction of the S8 state indicates durable depletion of a baseline-associated subset, while expansion of S6 reflects persistence of a distinct CD27^+^ population. Taken together, these findings show that B-cell remodeling after aHSCT involves sustained expansion of a single naïve state and reciprocal shifts within memory compartments, rather than simple restoration of baseline composition.

Although no states in the B_other compartment met the predefined criteria for persistent alteration (Fig. 4), its population-level trajectory was striking. B_other cells, which include CD20^low/−^ plasmablast/plasma-cell–like phenotypes and early transitional B-cell states, were profoundly depleted following conditioning and remained nearly absent through 12m before re-emerging at 24m (Fig. 3a). This delayed, non-linear recovery contrasts with the early expansion of naïve B cells and partial restoration of memory subsets. Together, these data show that B_other cells followed a markedly delayed reconstitution trajectory compared with conventional naïve and memory B-cell compartments.

### Summary of adaptive immune remodeling following aHSCT

Across adaptive lineages, aHSCT induced durable but asymmetric remodeling of immune architecture. In the B-cell compartment, reconstitution followed a hierarchical sequence: naïve B cells expanded early and remained elevated; memory B cells partially recovered with selective enrichment of CD27^+^ states and contraction of CD27^−^ subsets; and the B_other compartment, encompassing plasmablast/plasma-cell–like and transitional phenotypes, was profoundly depleted and re-emerged only at later time points. This temporal stratification indicates that B-cell reconstitution did not reflect uniform restoration of baseline distributions but instead followed distinct recovery dynamics across developmental stages. In contrast, T-cell remodeling was characterized by coordinated depletion of a discrete CD161^high^ chemokine receptor–rich module spanning CD8 memory and innate-like T-cell compartments, rather than staged developmental restructuring. Together, these findings show that adaptive reconstitution after aHSCT follows lineage-specific patterns, with hierarchical recovery dynamics in B cells and selective depletion of defined T-cell programs.

### Innate compartments show early redistribution with partial normalization by 24 months

Persistent state remodeling was confined to adaptive lineages. In contrast, innate compartments demonstrated largely transient redistribution without evidence of durable internal state reprogramming (eFigures 13 and 14). Within the NK compartment, CD16⁻ NK cells expanded early (100d) and remained elevated through 24m, whereas CD16⁺ NK cells declined from ∼6m and remained below BL and HC levels, indicating a sustained shift in NK subset composition. The monocyte and dendritic cell compartments showed early quantitative fluctuations, with partial normalization over time. The early predominance of CD16⁻ NK cells occurred concurrently with the contraction of CD161-enriched T-cell states, consistent with a transient regulatory milieu during immune reconstitution.

### Functional remodeling along a lineage-specific axis

To quantify functional reprogramming accompanying compositional remodeling, we projected single-cell marker profiles from BL and 24m into a supervised low-dimensional space optimized to maximize temporal separation (FreeViz). Within adaptive lineages that exhibited persistent architectural remodeling, BL and 24m populations occupied distinct regions of this functional remodeling space, indicating durable shifts in cellular phenotype beyond compositional changes (Fig. 6). Marker anchor distributions within this space suggested coordinated modulation of activation- and differentiation-associated programs across persistently remodeled states.

**Figure 6.**
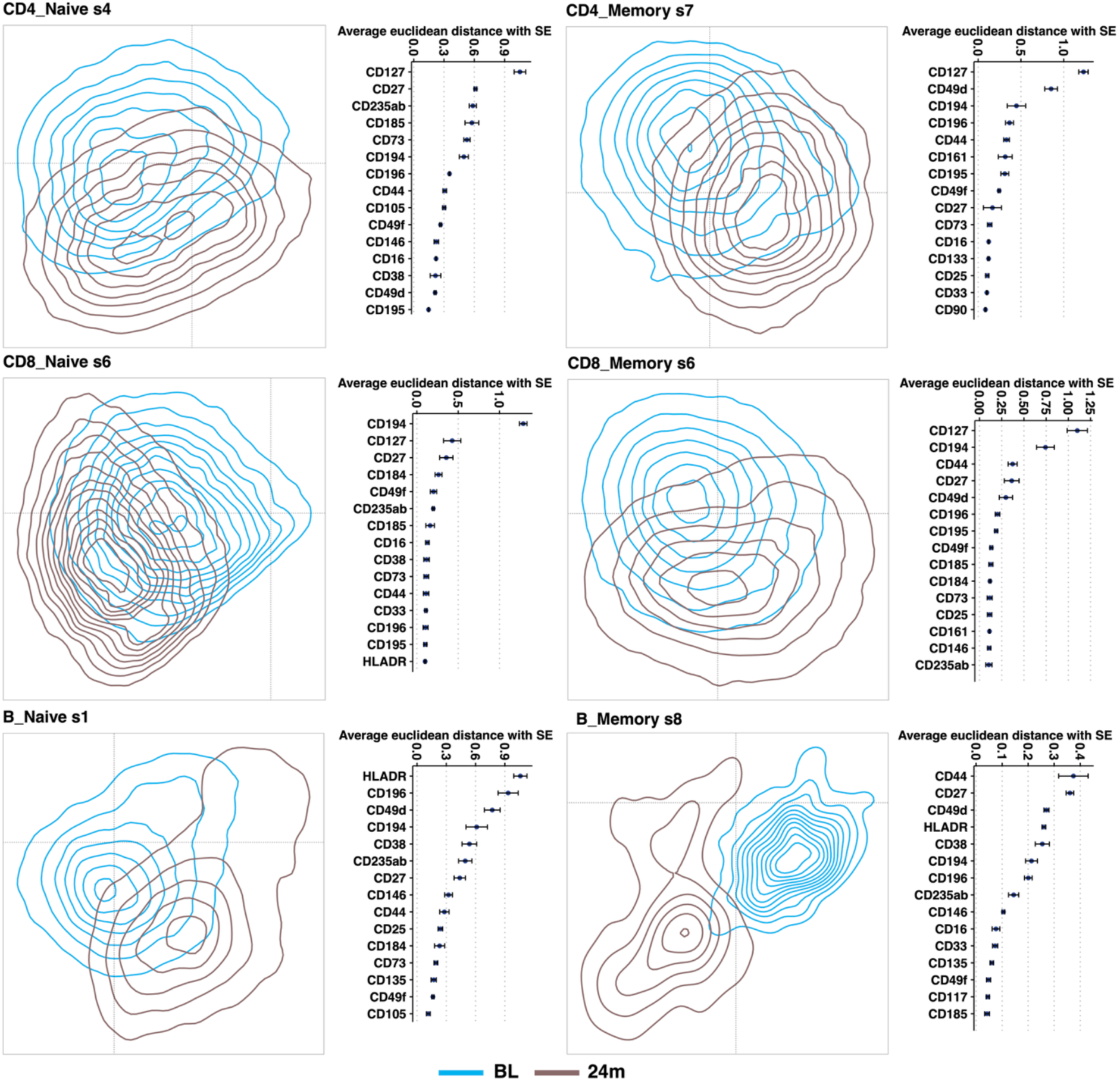
Functional separation space optimized for BL vs 24m discrimination. Two-dimensional density projections generated using the FreeViz algorithm comparing single-cell functional profiles at baseline (BL) and 24 months (24m) within adaptive immune compartments. In each of 1000 independent runs, equal numbers of randomly subsampled cells (n = 5000 per class, when available) were projected into a 2D space optimized to maximize separation between BL and 24m. Density contours represent aggregated projections across runs. Marker anchor positions indicate the relative contribution of individual functional markers to class separation. The mean Euclidean distance (± SE) of each marker from the origin is shown adjacent to each panel; markers positioned farther from the origin contribute more strongly to discrimination between time points. Distinct spatial segregation between BL and 24m populations reflects durable functional displacement along a lineage-specific remodeling axis following aHSCT.

Complementary state-level perturbation analysis using SARA supported this interpretation, demonstrating directional shifts in key differentiation and trafficking markers within cellular states exhibiting persistent abundance changes (eFigure 15). Although individual marker perturbations did not uniformly reach significance after multiple testing correction, the concordant directional patterns across states reinforce the presence of coordinated functional remodeling. Together, these analyses define a lineage-specific functional remodeling axis along which adaptive immune compartments undergo durable phenotypic displacement following aHSCT.

## Discussion

Autologous hematopoietic stem cell transplantation in treatment-refractory RRMS was associated with durable, lineage-specific remodeling of the adaptive immune system rather than simple restoration of the pre-treatment immune state. Although major immune populations partially normalized toward levels observed in healthy controls over 24 months, internal architectural reshaping within T- and B-cell compartments persisted, indicating that clinical remission occurred in the context of selective immune reorganization rather than a return to the pre-treatment immune state. Importantly, these effects were observed following lymphoablative rather than fully myeloablative conditioning, suggesting that profound adaptive immune remodeling can be achieved without complete myeloid ablation (26).

Within the T-cell compartment, naïve populations gradually repopulated, whereas memory CD4⁺ T cells remained reduced. Beyond changes in abundance, naïve T-cell reconstitution was accompanied by durable shifts in internal state composition, characterized by reduced representation of CD127-high homeostatic populations and expansion of CD127-negative and CD44⁺, CD38⁺ - or CD49d-expressing subsets. These findings indicate that naïve T-cell recovery reflects sustained compositional rebalancing rather than restoration of the pre-treatment state. In parallel, a coordinated CD161-expressing, chemokine receptor–enriched program spanning memory and innate-like subsets was persistently depleted. CD161⁺ T-cell populations have been linked to lineage commitment, CNS infiltration, and inflammatory activity in MS (27–31). Their sustained contraction across compartments therefore suggests selective depletion of pathogenic, inflammation-associated programs rather than indiscriminate lymphocyte loss. In the context of lymphoablative conditioning, these findings indicate contraction of specific pathogenic T-cell trajectories while permitting repopulation of other T-cell compartments.

Early after transplantation, we identified the emergence of a putative regulatory network that may establish a different immune responsive state. In line with previous studies of post-aHSCT reconstitution (16, 33), we observed a significant expansion of CD16⁻ NK cells. Given that NK cells can regulate Th17 responses implicated in MS pathogenesis (32), this early innate enrichment likely contributes to modulation of adaptive immunity during the critical reconstitution phase. Parallel to this innate shift, we observed an early expansion of T cells exhibiting a naïve regulatory phenotype (CD25^+^, CD127^-^, CD49d^-^). Together, these findings suggest that the early post-transplant period is defined by a coordinated, multi-lineage regulatory environment rather than simple numeric recovery.

In contrast, B-cell reconstitution followed a hierarchical developmental pattern. Naïve B cells expanded early and remained elevated, while memory B cells were durably reduced and selectively restructured. These findings indicate that B-cell reconstitution reflects sustained expansion of activation-associated naïve states together with reciprocal restructuring of memory compartments, rather than simple restoration of baseline distributions. The delayed reappearance of plasmablast and transitional-like populations suggests a deeper resetting of terminal differentiation programs compared with upstream naïve compartments. This interpretation is supported by the marked depletion and delayed reconstitution of the B_other compartment, indicating that terminal differentiation–associated programs are more profoundly disrupted than upstream naïve compartments. Given the established role of memory B-cell populations in MS pathophysiology and their therapeutic targeting in RRMS, this staged redevelopment may represent an important component of long-term immune stabilization.

The durable contraction of memory B-cell subsets observed after aHSCT resembles key immunologic features of anti-CD20 therapies, which target circulating memory B cells and are highly effective in RRMS (34). However, unlike the cyclical depletion–repopulation dynamics associated with repeated anti-CD20 dosing, aHSCT was associated with prolonged restructuring of B-cell maturation. Compared with monoclonal antibody–based immune reconstitution therapies such as alemtuzumab (4,11), aHSCT may establish a self-sustained adaptive immune configuration without ongoing immune suppression. Whether this structured reorganization contributes to differences in long-term durability or safety profiles warrants further investigation.

This study was conducted in a small cohort from a single treatment arm, which may limit generalizability. Immune profiling was restricted to peripheral blood and may not fully capture CNS compartment dynamics. Although longitudinal modeling identified persistent architectural remodeling, causal relationships between specific immune changes and clinical remission cannot be established. Functional validation and comparative analyses across immune reconstitution therapies will be important to further define mechanistic relevance.

Together, these findings indicate that adaptive immune architecture is durably reorganized rather than restored to its pre-treatment state along lineage-specific structural and functional axes following lymphoablation. Importantly, these changes occurred at the level of defined cellular states, indicating coordinated remodeling of immune architecture rather than uniform repopulation of naïve compartments. The asymmetry between contraction of inflammatory T-cell programs and restructuring of B-cell maturation indicates that aHSCT establishes a durably reconfigured adaptive immune landscape, involving coordinated remodeling of B-cell maturation and T-cell effector programs. Such selective remodeling provides a plausible immunologic substrate for the sustained clinical and radiological stability observed in this treatment-refractory RRMS cohort.

## Author contributions

This mass cytometry study was led and conceptualized by SG, and designed in collaboration with AKL, LB, ØT, AMO, EKK, JBH, NB, MB, KMM, and BTG. The paper was composed primarily by SG, DK, JBH, and YG. The RAM-MS trial was led by AKL, LB, ØT, AMO, and EKK. SG was responsible for the trial biobank SOP. Patient samples were collected and processed in Bergen by SG, JBH, SA, YG, and HO. Mass cytometry experiments were performed in Bergen by JBH, SG, SA, IH, YG, and HO. DK developed and wrote R scripts for mass cytometry data analyses. DK, SG, and JBH analyzed mass cytometry data. All authors contributed to data interpretation and writing. All authors read and approved the final manuscript.

## Conflicts of interest

SG, JBH, DK, YG, SA, IH, HO, NB, MB, BTG, AMO, EKK and AKL declare no conflicts of interest. KMM has received speaker honoraria from Biogen, Sanofi and Novartis; and honoraria for participation in scientific advisory board from Alexion, and has participated in clinical trials organized by Biogen, Merck, Novartis, Otivio, Sanofi and Roche. ØT has participated in advisory boards and received speaker honoraria from Biogen, Merck, Novartis, Teva, Roche, Sanofi, and Bristol Myers Squibb and has participated in clinical trials organized by Merck, Novartis, Roche, and Sanofi. LB has received unrestricted research grants to his institution and/or scientific advisory board or speakers’ honoraria from Biogen, Genzyme, Merck, Novartis, Roche, and Teva, and has participated in clinical trials organized by Biogen, Merck, Novartis, and Roche.

## Acknowledgments/funding

The authors gratefully acknowledge the Flow Cytometry Core Facility at the University of Bergen for their valuable technical support. We sincerely thank all patients who participated in the clinical trial, as well as their families, whose contributions have been essential for this study. We also extend our thanks to the dedicated laboratory personnel for their valuable assistance, particularly Hanne-Linda Nakkestad for her contribution throughout the project. Special thanks to Gerd Bringeland for optimizing part of the antibody panel used in this study. This work was supported financially by the Research Council of Norway, the Tom Wilhelmsen Foundation, the Odd Fellow MS foundation and Gerda Meyer Nyquist Guldbrandson og Gerdt Meyer Nyquists Legat.

## Notes

https://github.com/dkleftogi/immuneReconstitution_RRMS

### Competing Interest Statement

SG, JBH, DK, YG, SA, IH, HO, NB, MB, BTG, AMO, EKK and AKL declare no conflicts of interest. KM has received speaker honoraria from Biogen, Sanofi and Novartis; and honoraria for participation in scientific advisory board from Alexion, and has participated in clinical trials organized by Biogen, Merck, Novartis, Otivio, Sanofi and Roche. ØT has participated in advisory boards and received speaker honoraria from Biogen, Merck, Novartis, Teva, Roche, Sanofi, and Bristol Myers Squibb and has participated in clinical trials organized by Merck, Novartis, Roche, and Sanofi. LB has received unrestricted research grants to his institution and/or scientific advisory board or speakers honoraria from Biogen, Genzyme, Merck, Novartis, Roche, and Teva, and has participated in clinical trials organized by Biogen, Merck, Novartis, and Roche.

### Summary of Updates

Updated Table 1 to further anonymize patient demographic data by replacing exact ages with age ranges to preserve participant anonymity. Corresponding text describing baseline cohort characteristics was updated for consistency. No changes were made to the study design, analyses, results, figures, or conclusions.

