## Supplementary information for "Lineage-Specific Immune Remodeling Following Autologous Hematopoietic Stem Cell Transplantation in Relapsing–Remitting Multiple Sclerosis"

**eTable 1 – The antibody panel used for the mass cytometry analysis of samples.** Antibodies were selected based on our previous studies (1) and included markers for cell phenotype, functional chemokine receptors, adhesion molecules, and stem cell identification. Our panel is designed to investigate the role of cell trafficking in MS and immune reconstitution, which may help explain clinical recovery outcomes. In-house conjugations were done according to the manufacturer's Maxpar MCP9 and X8 labelling protocols.

| Label | Target | Antibody clone | Comment |
|---|---|---|---|
| 89Y | CD45 | HI30 | Leukocyte marker; Manual gating |
| 112Cd | CD4 | RPA-T4 | Annotation first round |
| 113Cd | CD14 | M5E2 | Annotation first/second round |
| 114Cd | CD19 | HIB19 | Annotation first round |
| 116Cd | CD3 | UCHT1 | Annotation first round |
| 141Pr | CD49d | 9F10 | Adhesion molecule; α4 integrin |
| 142Nd | CD73 | 606112 | Ectoenzyme purinergic signalling |
| 143Nd | HLA-DR | L243 | MHC class II |
| 144Nd | CD146 | P1H12 | Adhesion molecule MCAM |
| 145Nd | CD117 | 104D2 | Stem cell marker: c-Kit |
| 146Nd | CD8a | RPA-T8 | Annotation first round |
| 147Sm | CD20 | 2H7 | Annotation first/second round |
| 148Nd | CD34 | 581 | Hematopoietic stem cells |
| 149Sm | CD25 | 2A3 | Cytokine receptor (IL-2R) |
| 150Nd | CD105 | 166707 | Hematopoietic stem cells |
| 151Eu | CD278 | C398.4A | ICOS |
| 152Sm | CD66b | 80H3 | Neutrophil marker; Manual gating |
| 153Eu | CD194 | 205410 | Chemokine receptor (CCR4) |
| 154Sm | CD49f | MP4F10 | Adhesion molecule; integrin α6 |
| 155Gd | CD161 | HP-3G10 | KLRB1 |
| 156Gd | CD184 | 12G5 | Chemokine receptor (CXCR4) |
| 158Gd | CD27 | L128 | TNF receptor SF7; Annotation second round |
| 159Tb | CD45RO | UCHL1 | Annotation second round |
| 160Gd | CD44 | BJ18 | Adhesion molecule |
| 161Dy | CD235ab | HIR2 | Erythrocytes (bound to cells) |
| 162Dy | CD11c | Bu15 | Adhesion molecule; integrin αX; Annotation second round |
| 163Dy | CD33 | WM53 | Siglec-3 receptor; Annotation second round |
| 164Dy | CD133 | 170411 | Stem cell marker |
| 165Ho | CD127 | A019D5 | Cytokine receptor (IL7-Ra) |
| 166Er | CD123 | 6H6 | Cytokine receptor (IL-3R); Annotation second round |
| 167Er | CD162 | KPL-1 | Adhesion molecule |
| 168Er | CD185 | 51505 | Chemokine receptor (CXCR5) |
| 169Tm | CD90 | 5E10 | Stem cell marker |
| 170Er | CD45RA | HI100 | Annotation second round |
| 171Yb | CD195 | NP-6G4 | Chemokine receptor (CCR5) |

| | | | |
|---|---|---|---|
| 172Yb | CD38 | HIT2 | ADP ribose hydrolase enzyme |
| 173Yb | CD196 | G034E3 | Chemokine receptor (CCR6) |
| 174Yb | CD135 | 66903 | FLT3 |
| 175Lu | CD10 | HI10a | Neprilsyn enzyme |
| 176Yb | CD56 | HCD56 | Annotation first round |
| 209Bi | CD16 | 3G8 | FcγRIIIa; Annotation first/second round |

**eTable 2A – Major immune lineages.** Meta-clusters were manually inspected and annotated based on median marker expression profiles, resulting in identification of five major immune lineages:

| Lineage | Marker definition |
|---|---|
| CD4$^{+}$ T cells | CD3$^{+}$ CD4$^{+}$ CD8a$^{-}$ CD20$^{-}$ CD19$^{-}$ CD14$^{-}$ CD11c$^{-}$ CD33$^{-}$ CD56$^{-}$ |
| CD8$^{+}$ T cells | CD3$^{+}$ CD4$^{-}$ CD8a$^{+}$ CD20$^{-}$ CD19$^{-}$ CD14$^{-}$ CD11c$^{-}$ CD33$^{-}$ CD56$^{-}$ |
| B cells | CD3$^{-}$ CD4$^{-}$ CD8a$^{-}$ CD20$^{+}$ CD19$^{+}$ CD14$^{-}$ CD11c$^{-}$ CD33$^{-}$ CD56$^{-}$ |
| NK cells | CD3$^{-}$ CD4$^{-}$ CD8a$^{-}$ CD20$^{-}$ CD19$^{-}$ CD14$^{-}$ CD11c$^{-}$ CD33$^{-}$ CD56$^{+}$ |
| Myeloid / DCs | CD3$^{-}$ CD4$^{-}$ CD8a$^{-}$ CD20$^{-}$ CD19$^{-}$ CD14$^{+}$/$^{-}$ CD11c$^{+}$ CD33$^{+}$ |

**eTable 2B – Subpopulation annotations.** Each major lineage was subsequently re-clustered using the same FlowSOM and ConsensusClusterPlus strategy, followed by manual inspection and annotation. Seventeen subpopulations were identified.

| # | Parent lineage | Subpopulation | Marker definition |
|---|---|---|---|
| 1 | CD4$^{+}$ T | CD4$^{+}$ T memory | CD3$^{+}$ CD4$^{+}$ CD8a$^{-}$ CD45RA$^{-}$ CD45RO$^{+}$ |
| 2 | CD4$^{+}$ T | CD4$^{+}$ T naïve | CD3$^{+}$ CD4$^{+}$ CD8a$^{-}$ CD45RA$^{+}$ CD45RO$^{-}$ |
| 3 | CD8$^{+}$ T | CD8$^{+}$ T memory | CD3$^{+}$ CD4$^{-}$ CD8a$^{+}$ CD45RA$^{-}$ CD45RO$^{+}$ |
| 4 | CD8$^{+}$ T | CD8$^{+}$ T naïve | CD3$^{+}$ CD4$^{-}$ CD8a$^{+}$ CD45RA$^{+}$ CD45RO$^{-}$ |
| 5 | T cells | T_other | CD3$^{+}$ with ambiguous CD4/CD8a |
| 6 | T cells | T double-negative | CD4$^{-}$ CD8a$^{-}$ |
| 7 | B cells | B memory | CD20$^{+}$ CD19$^{+}$ CD27$^{+}$ |
| 8 | B cells | B naïve | CD20$^{+}$ CD19$^{+}$ CD27$^{-}$ |
| 9 | B cells | B_other | Ambiguous CD27/CD127 or high CD20/CD38 |
| 10 | NK cells | NK CD16$^{+}$ | CD56$^{+}$ CD16$^{+}$ |
| 11 | NK cells | NK CD16$^{-}$ | CD56$^{+}$ CD16$^{-}$ |
| 12 | NK cells | NK_other | Low CD56, ambiguous CD3/CD16 |
| 13 | Myeloid | Classical monocytes | CD14$^{+}$ CD16$^{-}$ CD33$^{+}$ |
| 14 | Myeloid | Non-classical monocytes | CD14$^{-}$ CD16$^{+}$ CD33$^{+}$ |
| 15 | Myeloid/DC | Conventional DCs | CD11c$^{+}$ CD123$^{-}$ CD14$^{-}$ CD16$^{-}$ CD33$^{+}$ |
| 16 | Myeloid/DC | Plasmacytoid DCs | CD11c$^{+}$ CD123$^{+}$ CD14$^{-}$ CD16$^{-}$ CD33$^{+}$ |
| 17 | Myeloid | Myeloid_other | Cells not confidently assigned |

**eTable 3 – Model-estimated $log_2$ fold changes from paired diffcyt-DA-edgeR analysis.** $Log_2$ fold changes ($log_2FC$) and FDR-adjusted p-values for each cell population at each post-aHSCT time point relative to baseline (BL), estimated using a paired negative binomial generalized linear model.

| CellType | Comparison | log2FC | FDR |
|---|---|---|---|
| CD4_Memory | 100d vs BL | -0.9 | 0.00 |
| CD4_Memory | 6m vs BL | -0.98 | 0.00 |
| CD4_Memory | 12m vs BL | -0.58 | 0.02 |
| CD4_Memory | 24m vs BL | -0.73 | 0.00 |
| CD4_Naive | 100d vs BL | -5.39 | 0.00 |
| CD4_Naive | 6m vs BL | -4.83 | 0.00 |
| CD4_Naive | 12m vs BL | -1.71 | 0.00 |
| CD4_Naive | 24m vs BL | -0.48 | 0.70 |
| CD8_Memory | 100d vs BL | 1.36 | 0.00 |
| CD8_Memory | 6m vs BL | 1.17 | 0.00 |
| CD8_Memory | 12m vs BL | 1.02 | 0.00 |
| CD8_Memory | 24m vs BL | 0.5 | 0.12 |
| CD8_Naive | 100d vs BL | -2.58 | 0.00 |
| CD8_Naive | 6m vs BL | -2.36 | 0.00 |
| CD8_Naive | 12m vs BL | -0.94 | 0.04 |
| CD8_Naive | 24m vs BL | 0.14 | 0.12 |
| T_DN | 100d vs BL | -1.36 | 0.00 |
| T_DN | 6m vs BL | -0.54 | 0.00 |
| T_DN | 12m vs BL | -0.25 | 0.16 |
| T_DN | 24m vs BL | 0.24 | 0.17 |
| T_other | 100d vs BL | 0.84 | 0.17 |
| T_other | 6m vs BL | -0.11 | 0.25 |
| T_other | 12m vs BL | 0.06 | 0.19 |
| T_other | 24m vs BL | 1.85 | 0.00 |
| B_Memory | 100d vs BL | -2.36 | 0.00 |
| B_Memory | 6m vs BL | -2.1 | 0.00 |
| B_Memory | 12m vs BL | -1.29 | 0.00 |
| B_Memory | 24m vs BL | -0.1 | 0.08 |
| B_Naive | 100d vs BL | 1.36 | 0.17 |
| B_Naive | 6m vs BL | 1.3 | 0.01 |
| B_Naive | 12m vs BL | 1.37 | 0.00 |
| B_Naive | 24m vs BL | 1.32 | 0.00 |
| B_other | 100d vs BL | 1.69 | 0.01 |
| B_other | 6m vs BL | 1.1 | 0.02 |
| B_other | 12m vs BL | 1.06 | 0.00 |
| B_other | 24m vs BL | 6.24 | 0.00 |
| NK_CD16_pos | 100d vs BL | 0.07 | 0.64 |
| NK_CD16_pos | 6m vs BL | 0.07 | 0.81 |

| | | | |
|---|---|---|---|
| NK_CD16_pos | 12m vs BL | -0.23 | 0.47 |
| NK_CD16_pos | 24m vs BL | -0.35 | 0.02 |
| NK_CD16_neg | 100d vs BL | 1.46 | 0.00 |
| NK_CD16_neg | 6m vs BL | 1.55 | 0.00 |
| NK_CD16_neg | 12m vs BL | 1.55 | 0.00 |
| NK_CD16_neg | 24m vs BL | 1.25 | 0.00 |
| NK_other | 100d vs BL | 0.51 | 0.29 |
| NK_other | 6m vs BL | 0.6 | 0.81 |
| NK_other | 12m vs BL | -0.77 | 0.95 |
| NK_other | 24m vs BL | 0.58 | 0.00 |
| Monocytes_classical | 100d vs BL | 0.15 | 0.17 |
| Monocytes_classical | 6m vs BL | 0.35 | 0.00 |
| Monocytes_classical | 12m vs BL | 0.06 | 0.47 |
| Monocytes_classical | 24m vs BL | -0.31 | 0.08 |
| Monocytes_nonClassical | 100d vs BL | 0.85 | 0.00 |
| Monocytes_nonClassical | 6m vs BL | 0.73 | 0.00 |
| Monocytes_nonClassical | 12m vs BL | 0.23 | 0.43 |
| Monocytes_nonClassical | 24m vs BL | -0.41 | 0.08 |
| DC | 100d vs BL | 0.03 | 0.27 |
| DC | 6m vs BL | 0.09 | 0.85 |
| DC | 12m vs BL | -0.2 | 0.42 |
| DC | 24m vs BL | -0.36 | 0.03 |
| DC_plasmocytoid | 100d vs BL | -0.36 | 0.23 |
| DC_plasmocytoid | 6m vs BL | -0.3 | 0.68 |
| DC_plasmocytoid | 12m vs BL | -0.17 | 0.97 |
| DC_plasmocytoid | 24m vs BL | -0.37 | 0.14 |
| Myelocytes_other | 100d vs BL | -0.6 | 0.06 |
| Myelocytes_other | 6m vs BL | 0 | 0.81 |
| Myelocytes_other | 12m vs BL | 0.11 | 0.47 |
| Myelocytes_other | 24m vs BL | 1.42 | 0.00 |

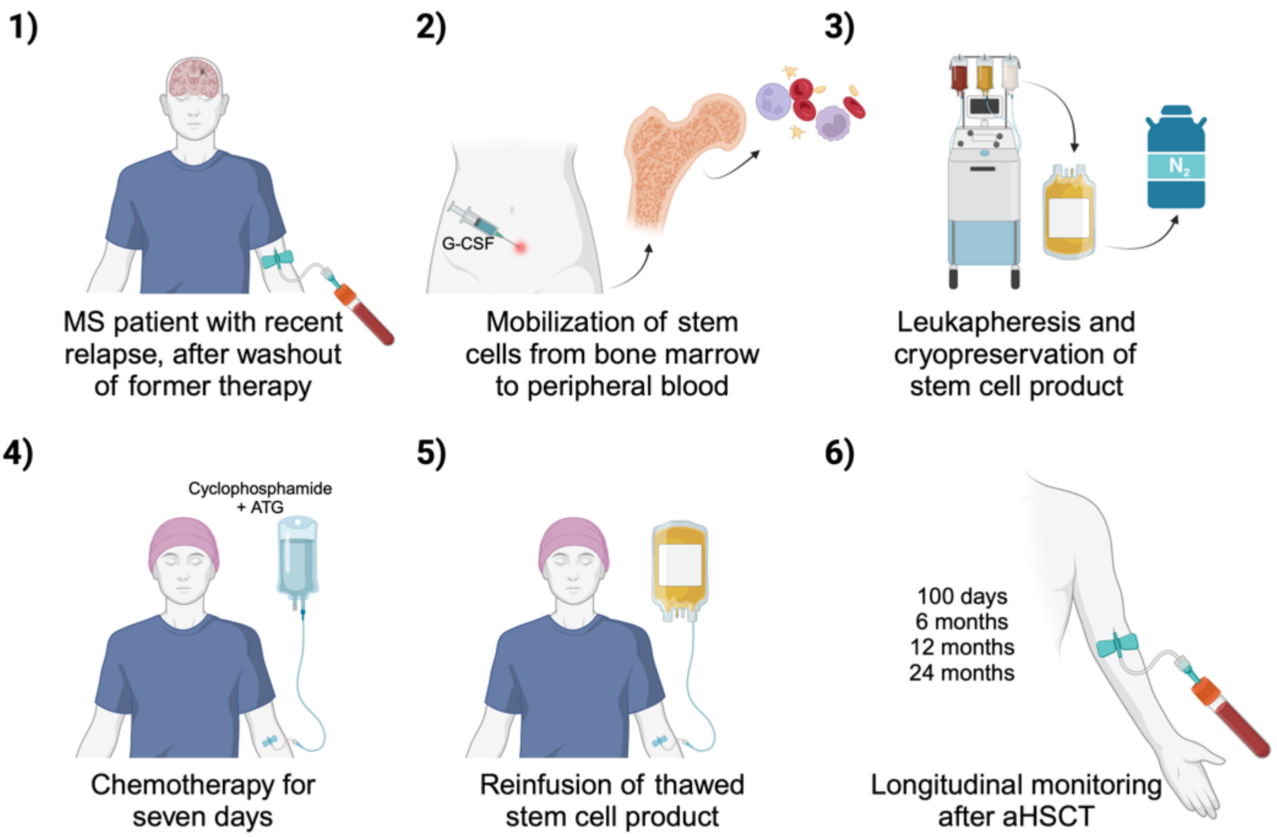


**eFigure 1 – aHSCT procedure in Arm A of the RAM-MS clinical trial.** The autologous hematopoietic stem cell transplantation (aHSCT) protocol for MS patients in Arm A consists of six key steps: **1)** Inclusion of patients following a recent relapse and therapy washout; **2)** Mobilization of stem cells to the peripheral blood; **3)** Collection of stem cells via leukapheresis, followed by cryopreservation; **4)** Administration of a seven-day chemotherapy regimen; **5)** Reinfusion of the thawed stem cell product; **6)** Longitudinal monitoring through peripheral blood sampling at 100 days, 6 months, 12 months, and 24 months post-transplantation.

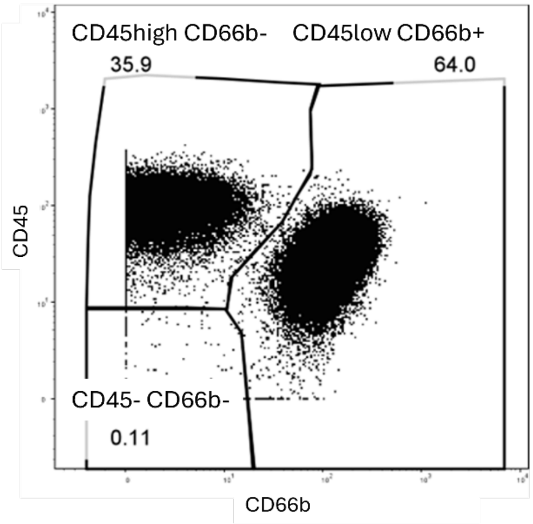


**eFigure 2 – Gating of single peripheral blood cells.** Three distinct populations are shown: CD45highCD66-, CD45lowCD66+, and CD45-CD66b-. The CD45highCD66b− gated cells (lymphocytes, monocytes, NK cells, dendritic cells) were further used for downstream analysis.

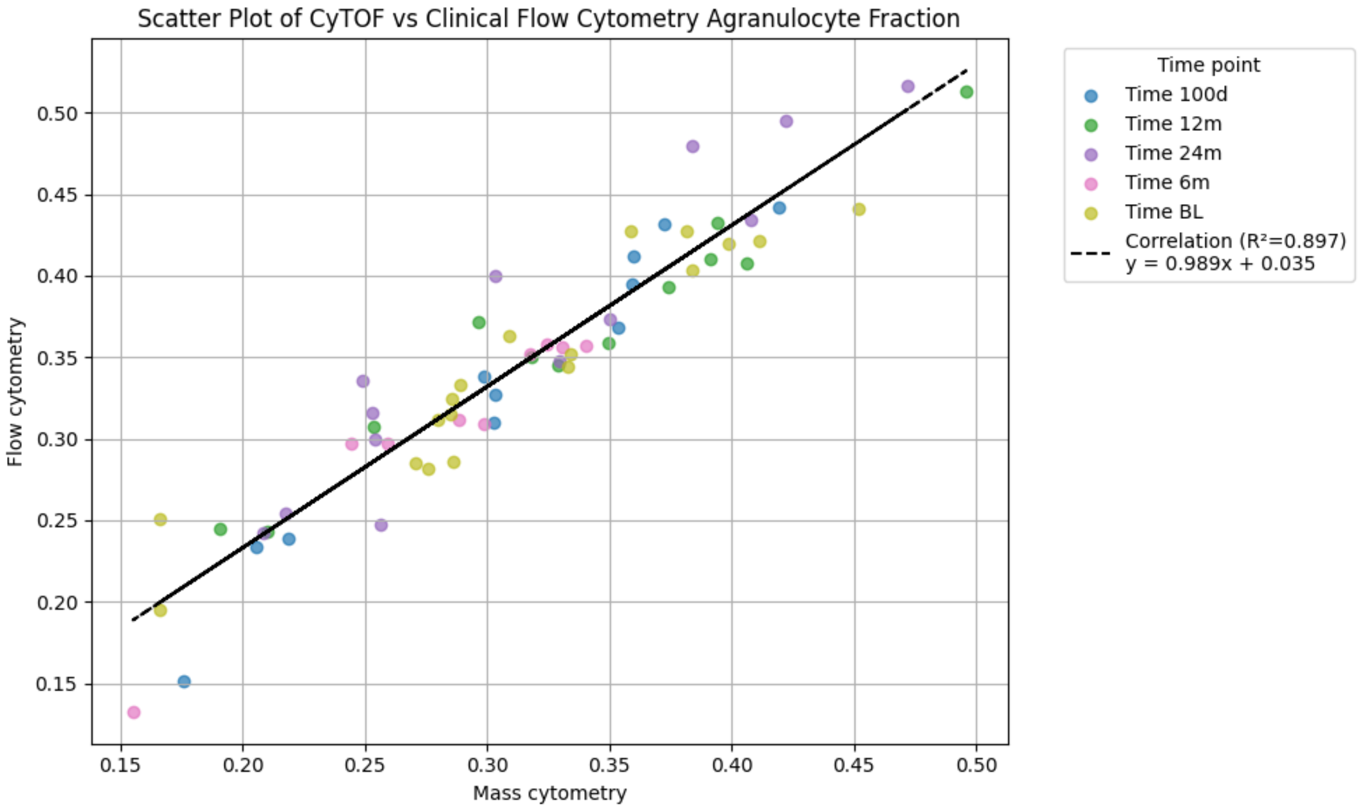


**eFigure 3 – Comparison between CyTOF and routine flow cytometry data.** Scatter plot showing the lymphocyte fraction in fixed PB samples analysed with mass cytometry (CyTOF), with the corresponding differential counts obtained in fresh whole blood analysed by conventional flow cytometry at the routine laboratory at Haukeland University Hospital. Using the conventional flow cytometry technique, the lymphocyte fraction is the sum of the lymphocyte and monocyte counts, divided by the total leukocyte count.

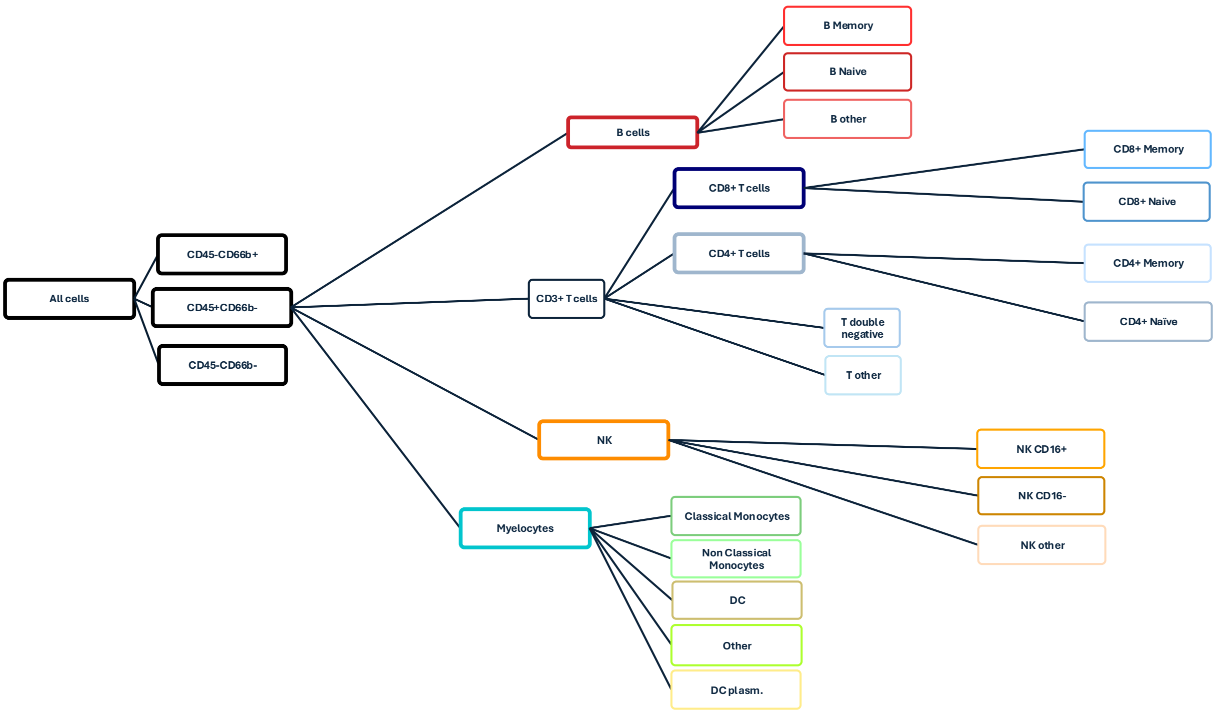


**eFigure 4 – Schematic of gating and clustering process into distinct cell populations.** We show an overview of the hierarchical scheme used to identify and name cell populations. Based on the manual gating shown in eFigure 2, the CD45 high CD66b- gate was used for downstream analysis. Cells in this gate were subjected to two rounds of clustering and meta-clustering. In the first round, five lineage subpopulations were detected. In the second round, each subpopulation was clustered/meta-clustered independently, resulting in 17 cell populations/clusters, as shown in the rightmost part of the figure.

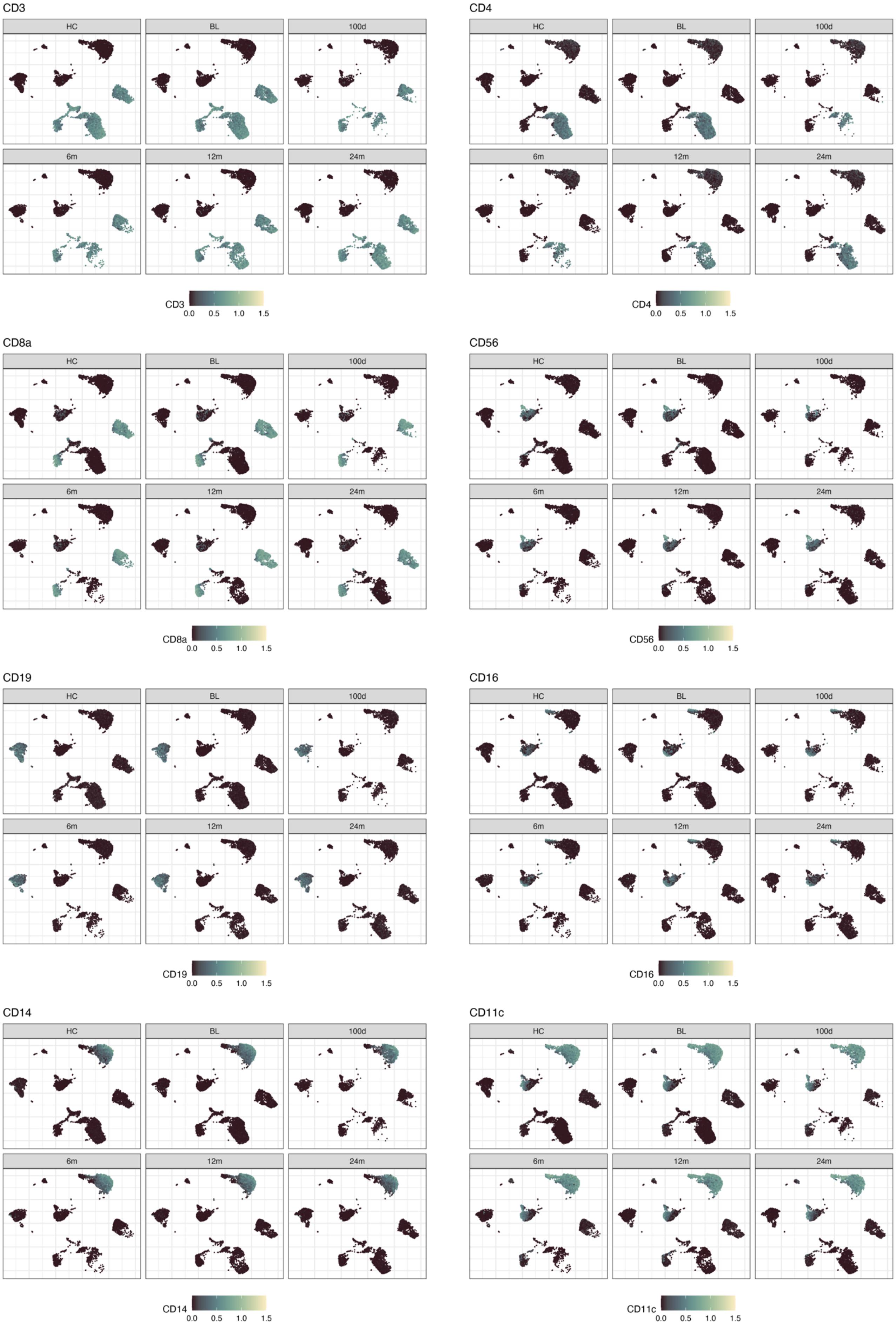


**eFigure 5 – UMAP visualisation of immune marker expression.** UMAP plots showing a randomly selected subset of 50,000 cells, with the expression of canonical immune markers overlaid at baseline (BL), 100 days (100d), 6 months (6m), 12 months (12m), 24 months (24m), and in healthy controls (HC).

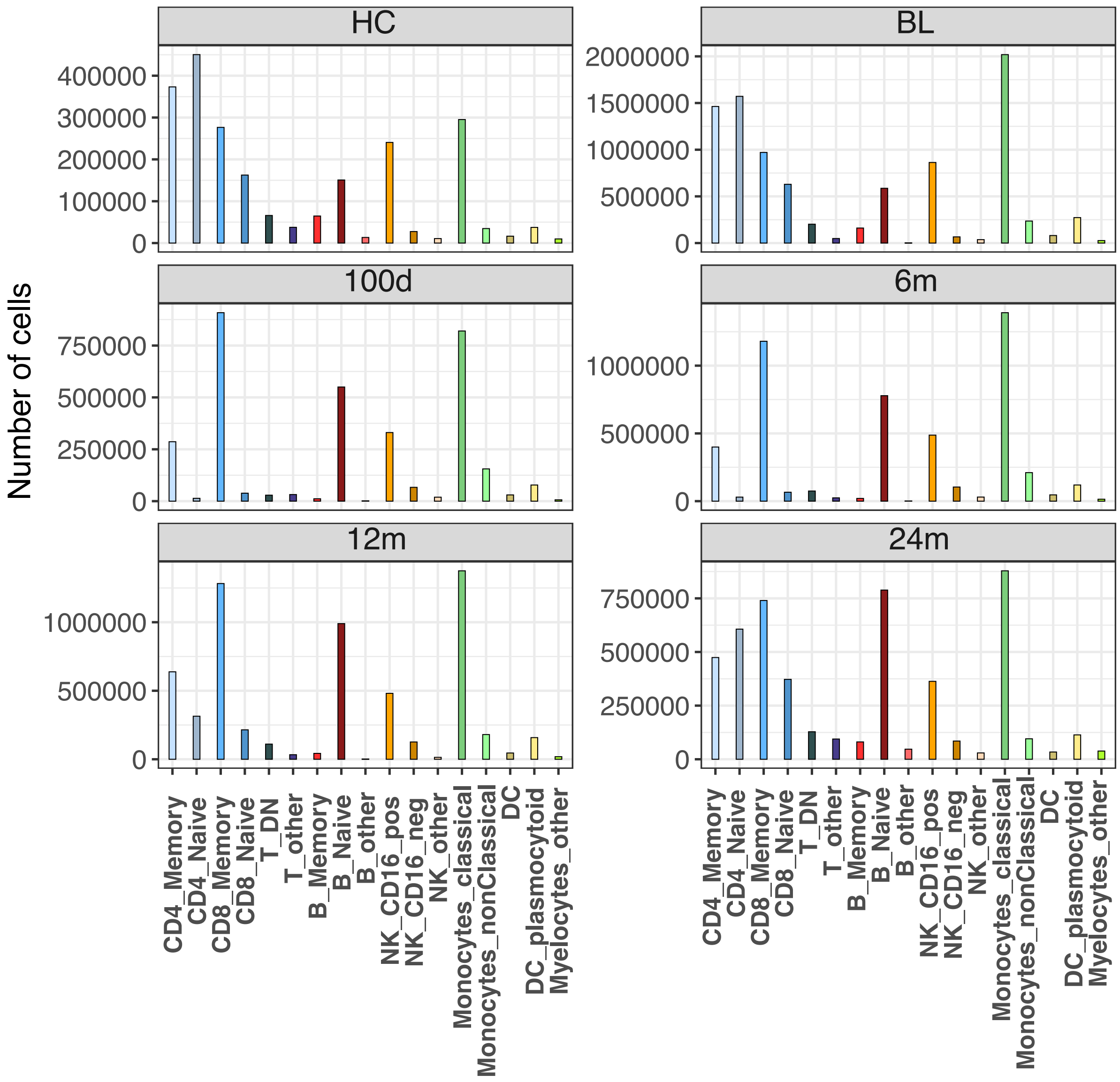


**eFigure 6 – Number of cells analysed per population across time points.** The number of cells analysed in each indicated immune cell population at baseline (BL), 100 days (100d), 6 months (6m), 12 months (12m), and 24 months (24m), as well as in healthy controls (HC), is shown. Cell counts are colour-coded for each time point.

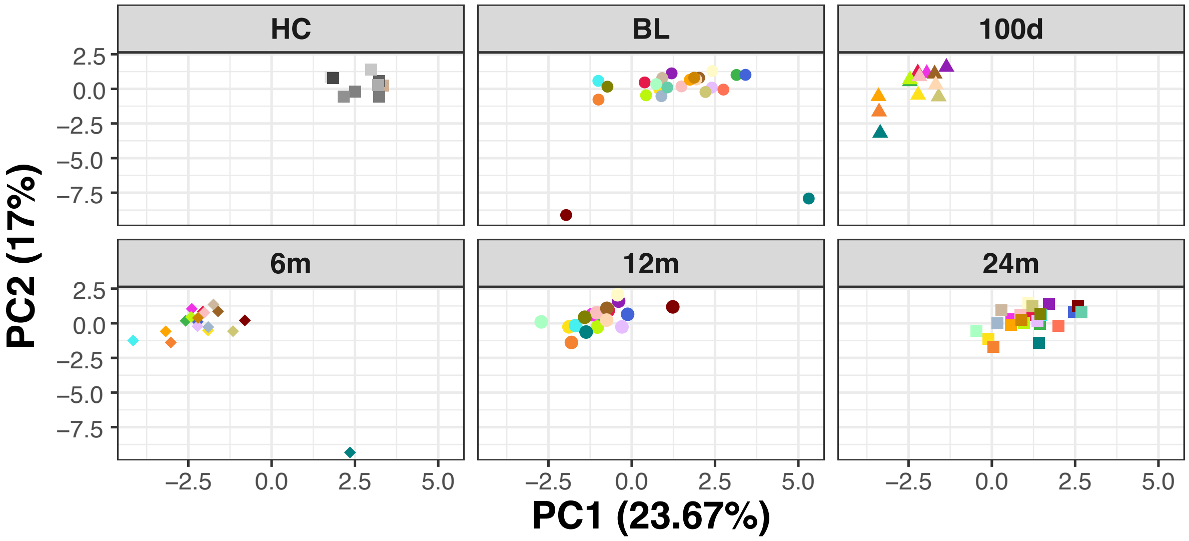


**eFigure 7 – Principal Component Analysis (PCA) of immune cell populations.** PCA was performed using the relative abundance of the 17 immune cell clusters described in Figure 2. Each colour represents an individual patient at the indicated time points: baseline (BL), 100 days (100d), 6 months (6m), 12 months (12m), and 24 months (24m). Healthy controls (HC) are shown in grey.

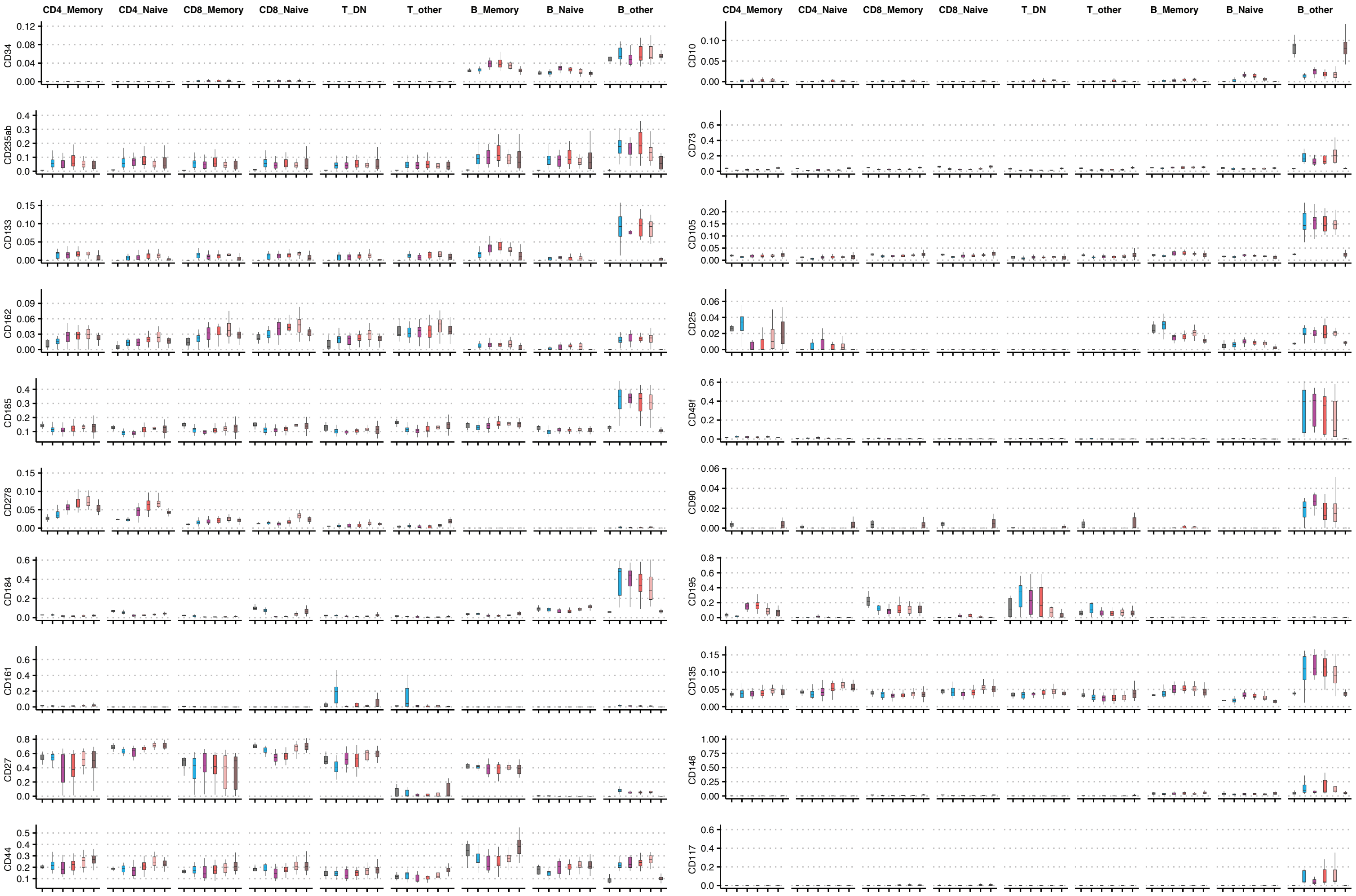


**eFigure 8 – Expression of markers across T and B cell populations.** Boxplots depicting median marker expression levels across all patients. CD34, a marker expressed on hematopoietic stem cells and gradually lost during hematopoiesis, was expressed in B cell clusters but not T cell clusters, where the expression increased post-aHSCT and thereafter returned to BL and HC levels. CD235ab was systemically expressed in both the T cell and B cell compartments in RRMS patients but not in HC. Similar to observations in systemic lupus erythematosus (SLE), a systemic autoimmune disease, erythrocytes may attach to lymphocytes due to high expression of adhesion molecules in RRMS patients. The highest expression was observed in the B_other cluster. CD162 (PSGL-1) showed an increasing trend across the T and B cell compartments post-aHSCT, whereas CD49f (integrin α6) expression decreased across T cell compartments but remained unchanged in B cells. CD184 (CXCR4) expression declined post-aHSCT. CD195 (CCR5) expression transiently increased between 100 days and 6 months post-aHSCT in naïve and memory CD4 cells and naïve CD8 T cells, while it decreased in memory CD8 T cells, T_DN and T_other cells. B cells did not express CD195. CD185 (CXCR5) expression increased between 6 and 12 months post-transplantation. CD146 (MCAM) transiently decreased in the B_other cluster, while CD135 (FLT3) levels increased in naïve CD4 T cells and naïve B cells but remained relatively stable in memory T cells. Lastly, CD278 (ICOS) expression increased in T cells post-aHSCT, and CD44 expression increased in naïve CD4 T, CD8 T cells and naïve B cells, but remained relatively stable in the memory compartments.

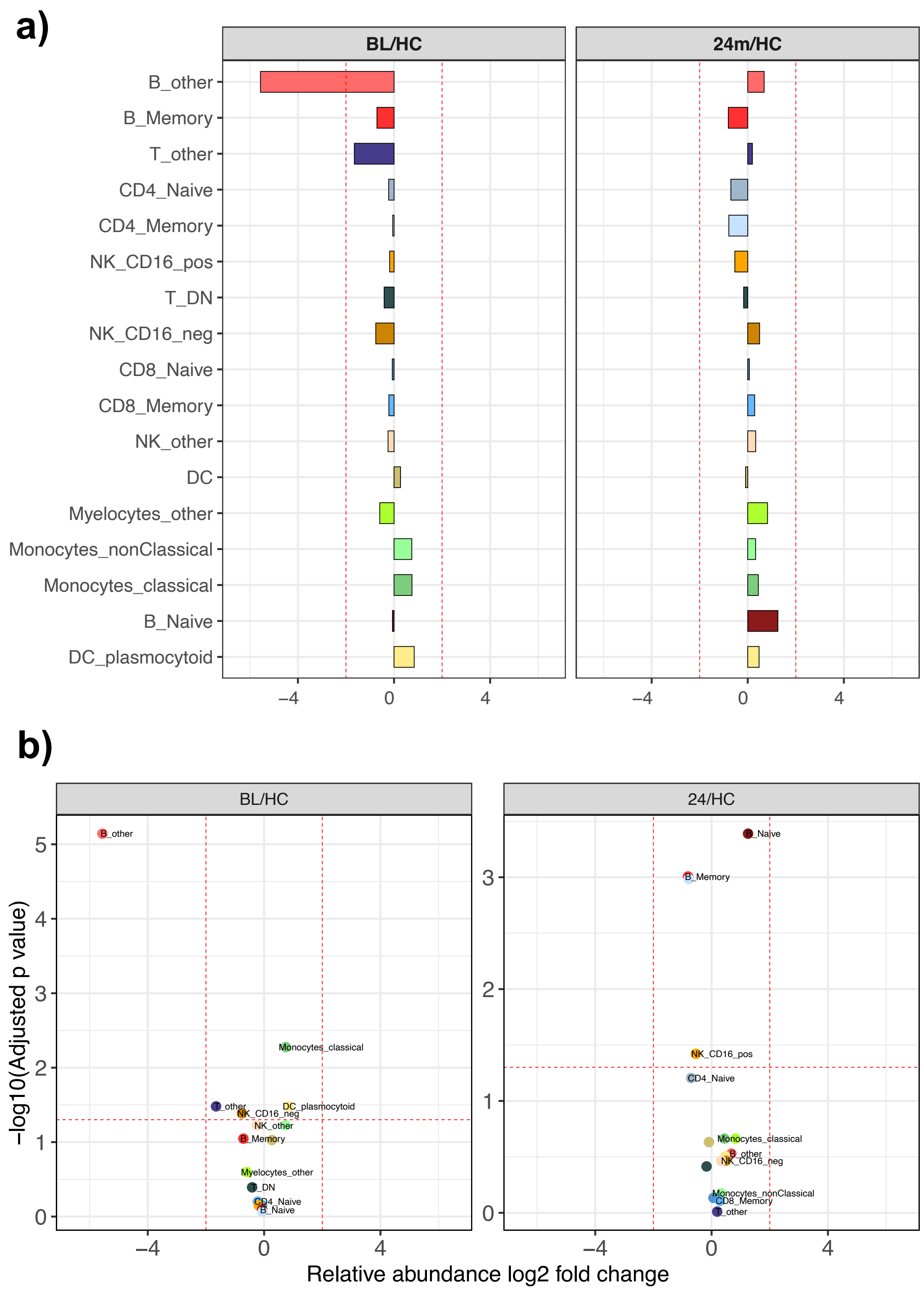


**eFigure 9 – Comparison of cell populations to healthy controls. a,** Bar plots showing log2 fold changes (log2 FC) for patient samples at baseline and at 24 months, relative to healthy controls (HC). Cell populations are ordered from highest to lowest average log2 FC; **b,** Volcano plots summarising differential abundance analysis performed using the diffcyt package, applying a p-value cutoff of 0.05 after multiple comparison correction and log2 FC thresholds of ±2.

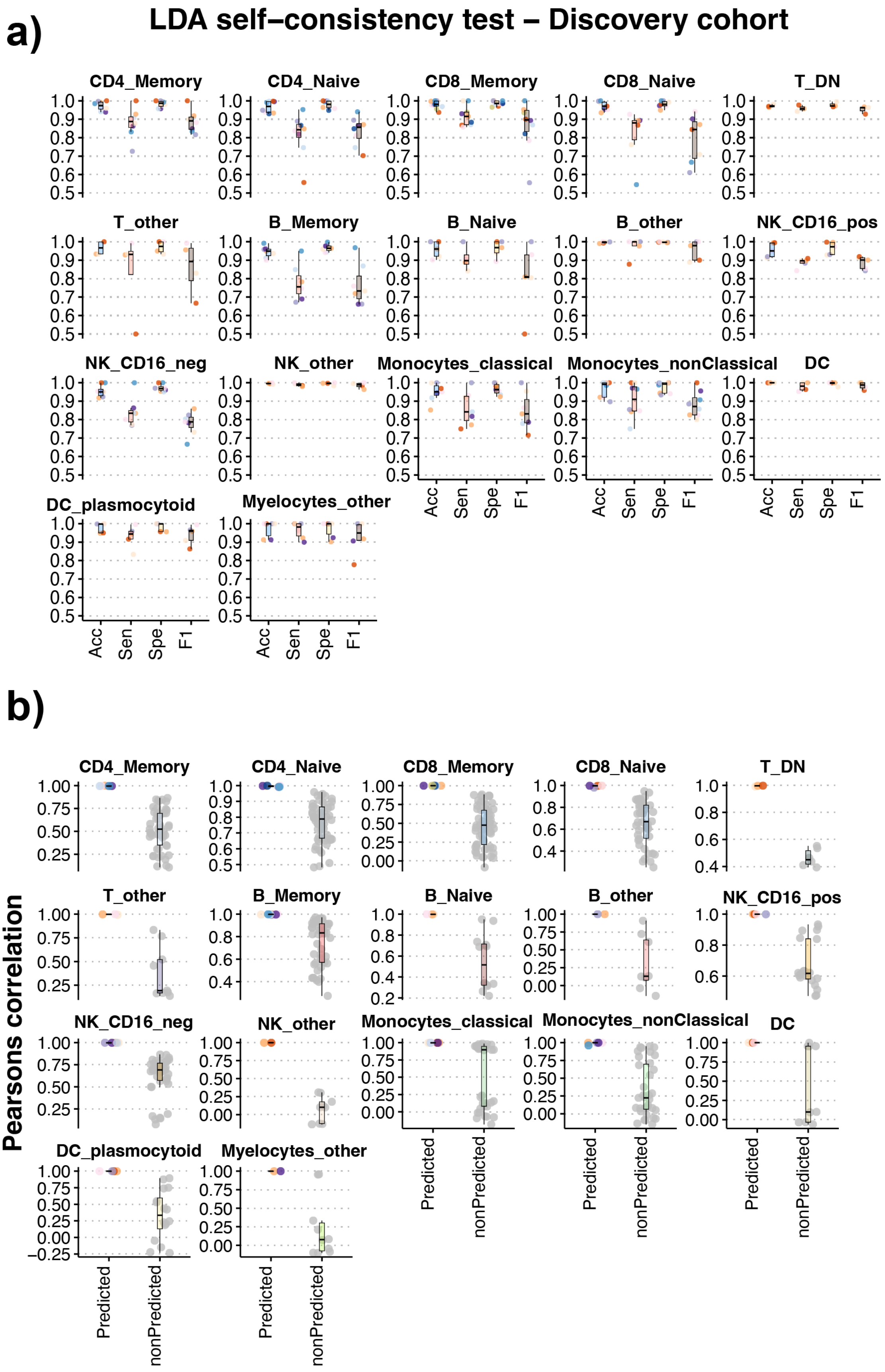


**eFigure 10 – Performance assessment of cellular state identification using LDA in the discovery cohort. a**, Performance of the cell type-specific models developed for cellular state identification. Each subplot shows the performance metrics for one model, evaluated in a hold-out setting using Accuracy (Acc), Sensitivity (Sen), Specificity (Spe), and F1 score; **b,** Pearson correlation analysis between the expression profiles of the correctly predicted cellular states (Predicted) and all other cellular states (non-Predicted).

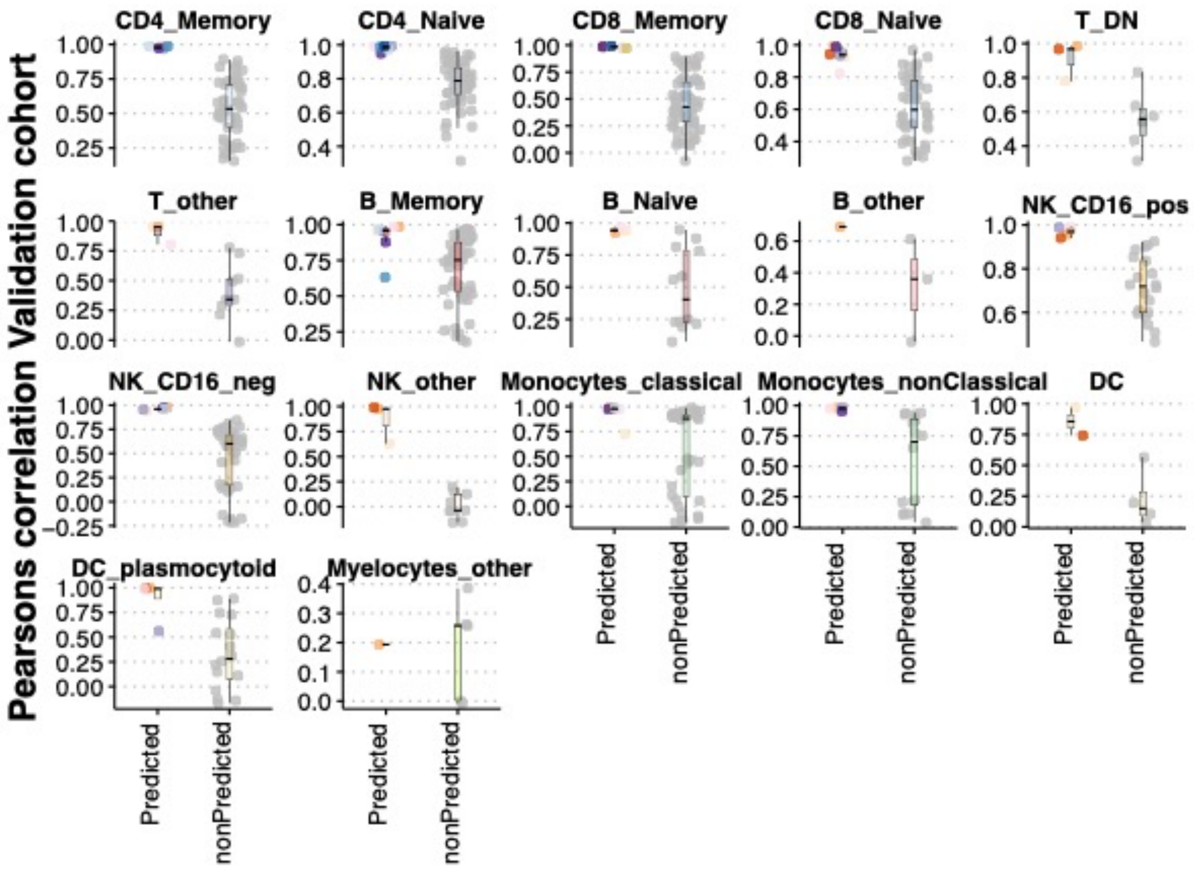


**eFigure 11 – Performance assessment of cellular state identification using LDA in the temporal extension cohort.** Each subplot illustrates the Pearson correlation between the expression profiles of the predicted state and all other cellular states found in each cell population.

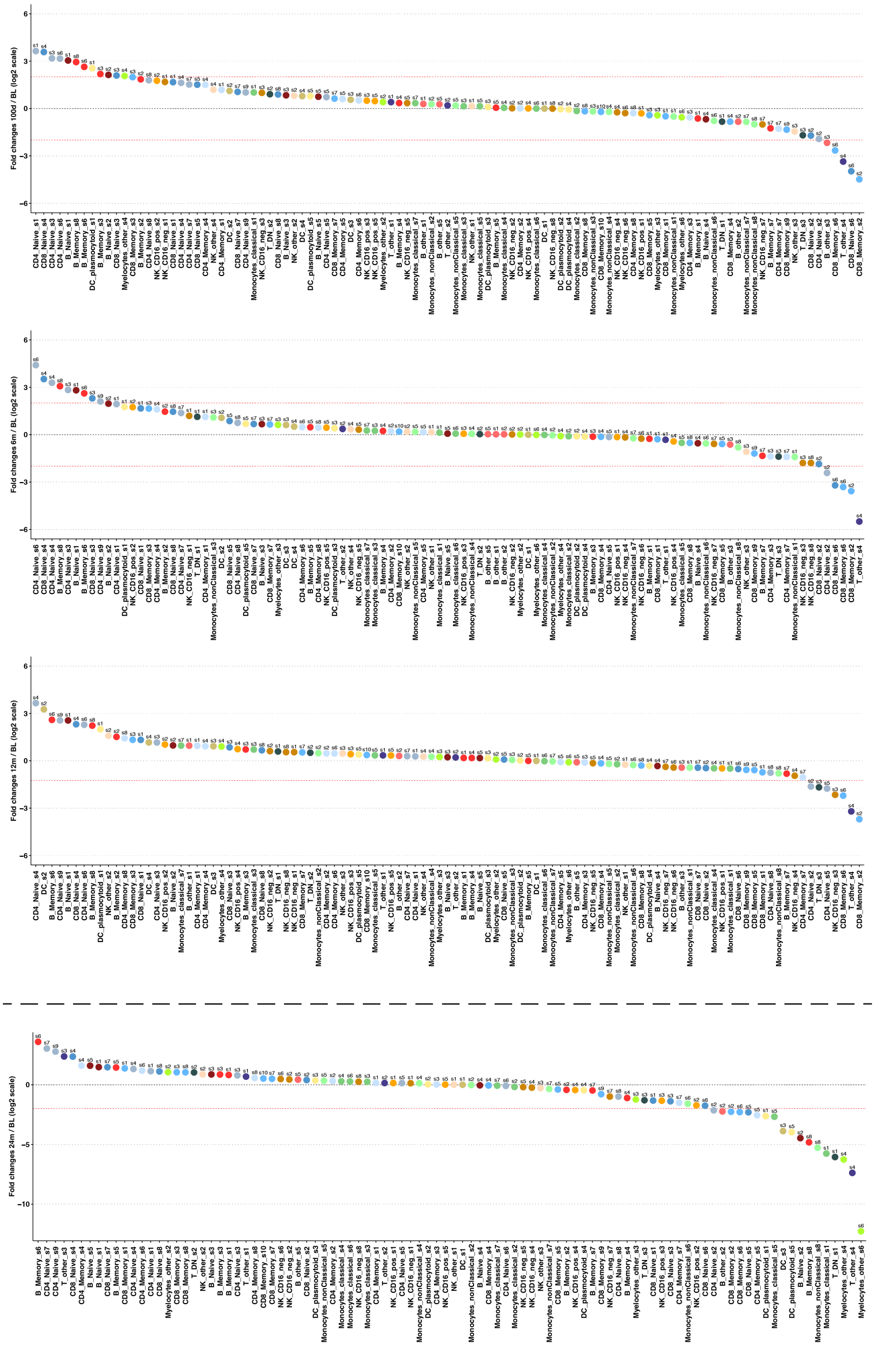


**eFigure 12 – Cellular state relative abundance.** Dot plots showing log2 fold changes from baseline (BL) to 100 days (100d), 6 months (6m), 12 months (12m), and 24 months (24m, temporal extension cohort) for each cellular state identified in the discovery cohort. The dot plots are ordered from the highest log2 FC to the lowest. The cell states are numbered and colour-coded based on their respective cell population.

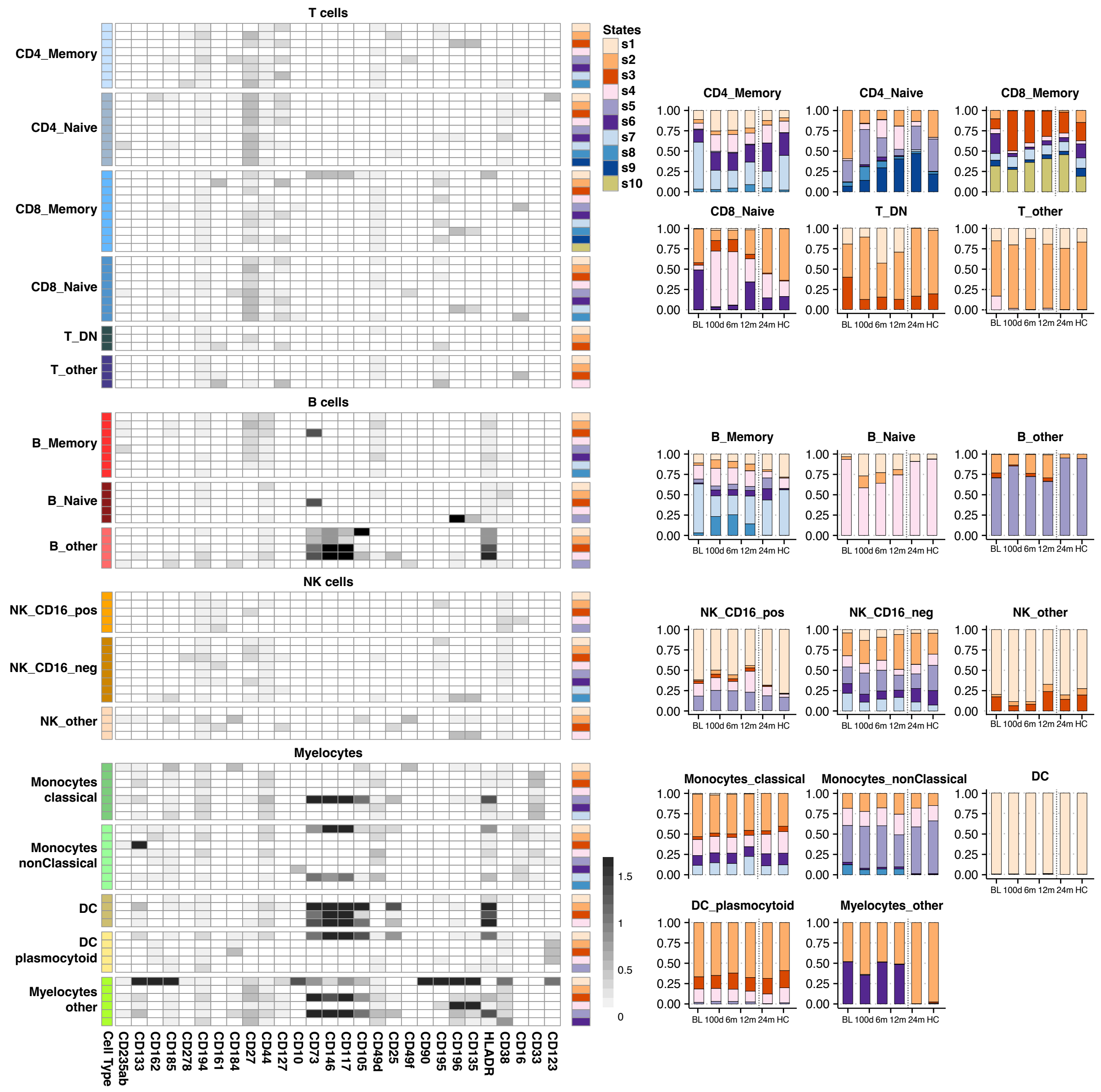


**eFigure 13 – Functional cellular states identified and longitudinally mapped across immune populations.** Left: Heatmap showing average expression profiles of 107 cellular states defined by unsupervised clustering within the discovery cohort (BL-12m), based on 28 additional markers. Right: Relative abundance of these predefined cellular states across all timepoints, including projected states for the temporal extension cohort (24m and HCs), assigned using the trained LDA models. States are displayed within their respective lineage-defined immune populations. This unified representation allows direct biological comparison of functional state dynamics before and after aHSCT, and relative to healthy controls.

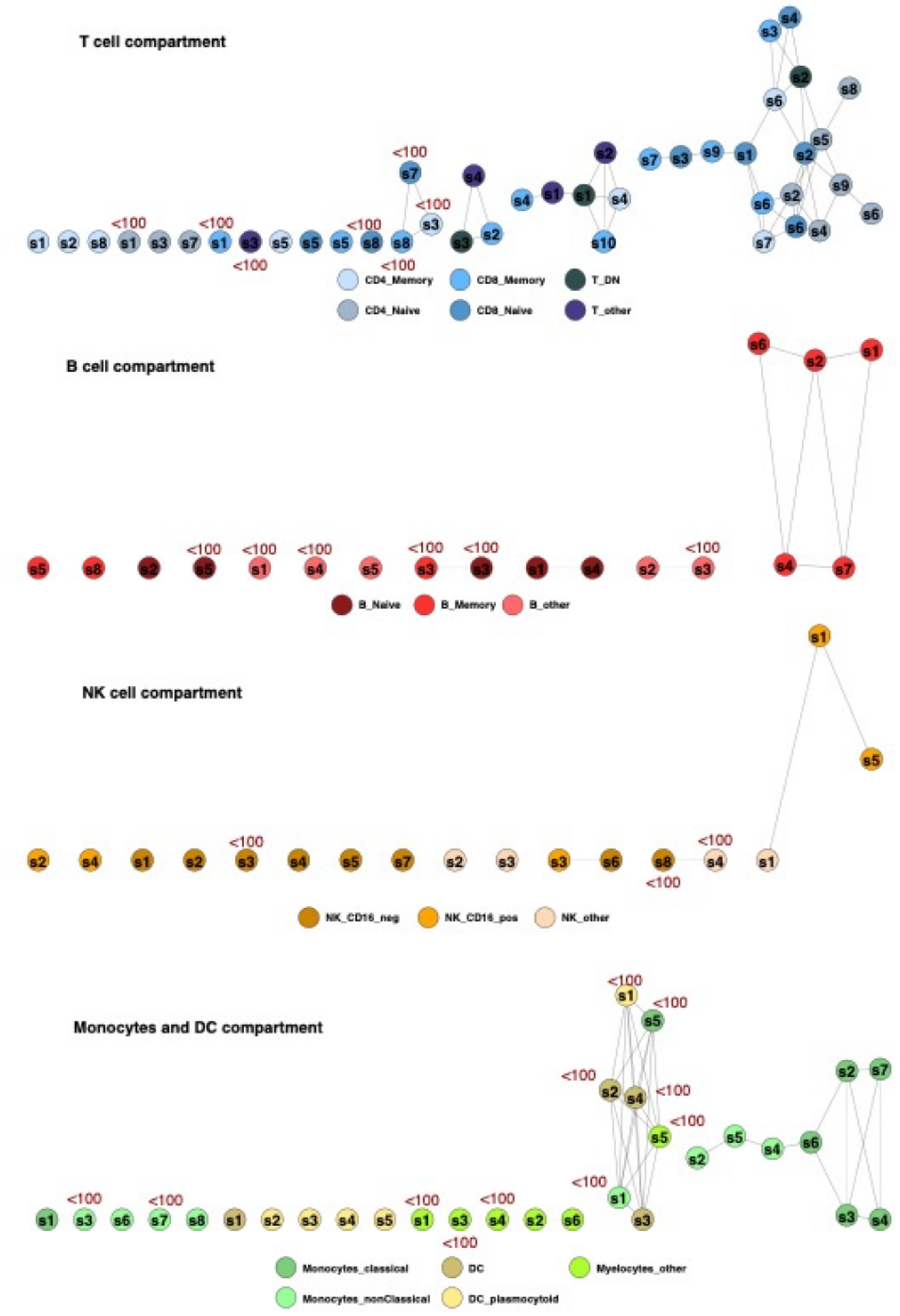


**eFigure 14 – Functional graphs of cellular state associations within each cellular compartment.** Graphs showing associations between the expression profiles of cellular states within each cellular compartment. Cellular states are connected by lines if their Pearson correlation exceeds 0.9. States containing fewer than 100 cells are specifically marked.

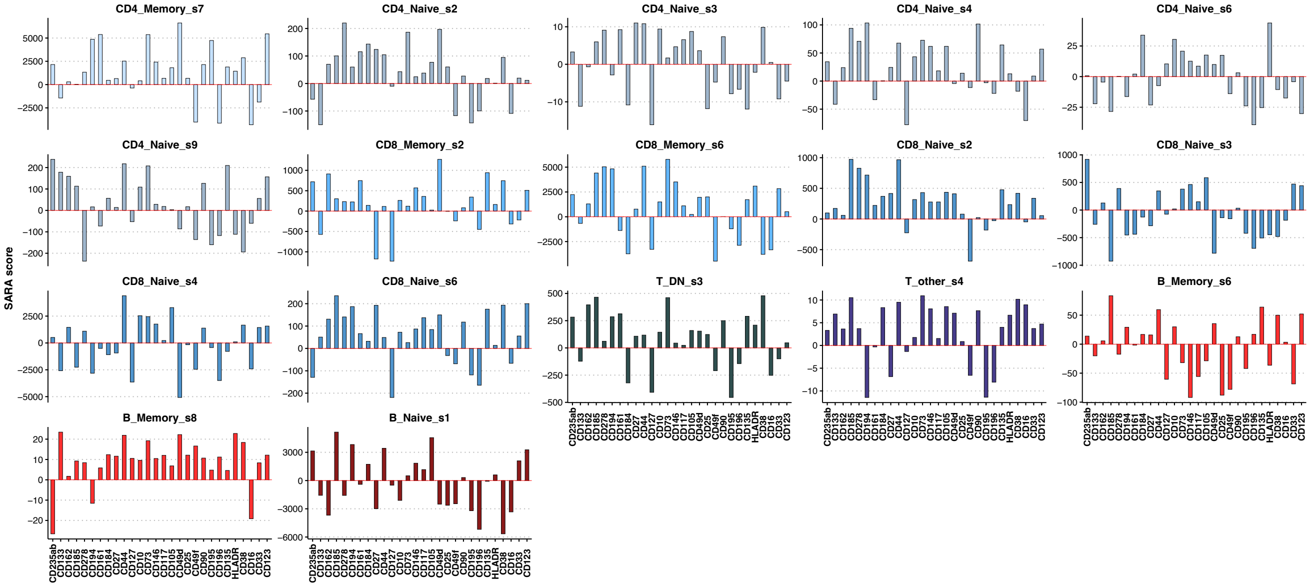


**eFigure 15 – Marker perturbation analysis using SARA.** For each functional marker, SARA scores compare single-cell intensity distributions between baseline (BL) and 24 months (24m) within cellular states that showed persistent changes in relative abundance. Positive scores indicate marker induction at 24m, whereas negative scores indicate inhibition. Scores are computed independently per cellular state and are therefore not directly comparable in magnitude across states.

**eMethods**

**Study Design and Cohort Characteristics**

This study included 25 patients with RRMS who received aHSCT in Arm A of the RAM-MS of the Randomized Autologous heMatopoietic Stem Cell Transplantation Versus Alemtuzumab, Cladribine or Ocrelizumab for RRMS (RAM-MS) clinical trial (NCT03477500; EudraCT 2014-510630-40-00). Patients were enrolled due to recent disease activity, despite receiving disease-modifying therapies, natalizumab, dimethyl fumarate, teriflunomide, fingolimod, interferon beta, and glatiramer acetate. The conditioning regimen consisted of lymphoablative therapy with cyclophosphamide total dose 200 mg/kg body weight given over 4 days and antithymocyte globulin (ATG) total dose 6 mg/kg body weight given over 5 days following stem cell mobilization with cyclophosphamide 2g/m2 body surface day 1 and granulocyte colony-stimulating factor (G-CSF) 5 micrograms/kg body weight daily for 5 days and 5 micrograms/kg body weight x 2 the day before the first harvesting day and until finalized harvesting.

The mean patient age was 38 years (16 females and 9 males). None of the patients had received prior T- or B-cell–depleting therapies, nor experienced clinical relapses during the drug-specific washout period defined by the RAM-MS protocol (ClinicalTrials.gov: NCT03477500), that would be expected to confound baseline immune phenotypes (Table 1).

Biobanked peripheral whole-blood samples obtained at baseline and 100 days, 6 months, 12 months, and 24 months post-aHSCT were used for immune profiling. A sex- and age-matched cohort of nine healthy individuals (5 females, 4 males, mean age 38) served as controls (eFigure 1).

The study was conducted in accordance with the approval of the Regional Committee for Medical and Health Research Ethics, Western Norway (REK 606396), and all individuals provided written informed consent.

**Mass cytometry and data acquisition**

Heparinized, biobanked 1 ml whole-blood samples stored at −80°C in Proteomic Stabilizer I (SMART TUBE Inc., San Carlos, CA, USA) were processed for mass cytometry analysis using standardized protocols for red blood cell lysis. Cells were counted, inspected, and 3 × $10^6$ cells were aliquoted in PBS and stored at −80°C. Samples were then barcoded using the Cell-ID™ 20-Plex Pd Barcoding Kit (Standard BioTools) [e1], pooled and stained with a previously prepared master mix of a 41-parameter mass cytometry antibody panel to identify innate and adaptive immune cell subsets (eTable 1). In-house-conjugated or commercially obtained metal-conjugated antibodies were titrated on pooled titration-samples combining mesenchymal stem cells, bone marrow cells, PBMCs, and buffy coats. Optimal concentrations were selected to minimise spillover according to a previously published protocol.

Data were acquired on a CyTOF XT (Standard BioTools) following our previously published protocols [e2]. Longitudinal samples (baseline to 12 months) from each patient were processed together and acquired within the same barcode to minimize intra-individual technical variation. Each barcode pool included a unique healthy donor reference sample to monitor batch-to-batch variability and enable inter-run batch correction [e3]. Twenty barcoded samples (Cell-ID™ 20 Plex-Pd Barcoding Kit (Standard Biotools) were pooled and stained with an aliquot of a previously prepared and frozen (–80°C ) antibody cocktail master mix. Thereafter, barcodes were stained overnight with iridium DNA intercalator at 4°C. The following day, barcoded samples were washed three times with metal-free cell staining buffer, resuspended in metal-free cell acquisition buffer, filtered (35μm), pelleted into 7-12 acquisition tubes, and loaded into the autosampler carousel of the CyTOF XT. Events were acquired at 300–500 events/sec using Acquisition Solution (CAS) Plus (Standard Biotools), with event rates monitored and adjusted throughout the 22-24-hour acquisition run. EQ Six Element Calibration Beads (Standard BioTools, PN 201245) and CAS were loaded into the autosampler carousel and added to each tube prior to acquisition for within-run signal normalization. In total, eleven 20-sample barcode pools (baseline–12 months) were stained and acquired over sequential 22–24–hour runs, resulting in 13 consecutive days of CyTOF XT acquisition with over 200 million total events. Prior to each 22–24-hour barcode acquisition run, the CyTOF XT instrument was tuned, cleaned, and maintained according to the manufacturer's recommended procedures to ensure stable signal performance across runs.

Twenty-four–month samples and age- and sex-matched healthy controls were acquired approximately one year later on the same CyTOF XT instrument using identical antibody master mixes, barcoding strategy, calibration beads, healthy reference controls, and acquisition settings. The same normalization and batch-correction pipeline was applied across both acquisition periods to ensure comparability of longitudinal patient samples and healthy controls.

**Mass cytometry data processing and cell population identification**

Each barcoded pool was normalized to EQ Six Element Calibration Beads and debarcoded with the R package Premessa (https://github.com/ParkerICI/premessa) with file-specific separation cutoffs. The debarcoded samples were then exported as individual FCS files, inspected, EQ beads excluded, and singlets identified based on 191Ir/193Ir DNA intercalators intensity and event length, using Cytobank and FlowJo software. The manually gated CD45$^{high}$CD66b$^{-}$ events (eFigure 2) were used for downstream analysis. The dataset was transformed using the arcsinh transformation, with marker-specific cofactors applied to stabilize variance and account for signal intensity differences across runs and barcodes. To minimize residual inter-run batch effects, marker intensities were aligned across barcode-specific healthy reference controls by cofactor adjustment to achieve comparable mean labeling intensities across the controls of all barcodes, following principles described by Ingelfinger et al. [e4]. Concordance between CyTOF-derived fractions and routine clinical flow cytometry differentials was assessed as an external QC step (eFigure 3).

Two sequential rounds of clustering and meta-clustering were implemented hierarchically using FlowSOM (10x10 grid) [e5] followed by ConsensusClusterPlus [e6] to identify major cell populations within the CD45$^{high}$CD66b$^{neg.}$ gate. Clusters were defined based on positive lineage-defining markers and the absence of alternative lineage signatures; marker combinations are shown in full in eTable 2. Eighteen antibodies from the 41-antibody panel were selected for clustering based on lineage and subset discrimination capacity: CD3, CD4, CD8a, CD45RA, CD45RO, CD20, CD19, CD27, CD117, CD56, CD127, CD34, CD38, CD16, CD14, CD11c, CD123, and CD33.

Meta-clusters were manually inspected and annotated based on median marker expression profiles, resulting in five major immune lineages: CD4+ T cells (CD3+CD4+); CD8+ T cells (CD3+CD8a+); B cells (CD19+CD20+); NK cells (CD3−CD56+); and Myeloid and dendritic cells (CD14/CD11c/CD33+). Clusters were defined based on positive lineage-defining markers and absence of alternative lineage signatures, as confirmed by median marker expression heatmaps.

Each major immune lineage was subsequently re-clustered using the same FlowSOM and ConsensusClusterPlus strategy, followed by manual inspection and annotation. This hierarchical approach identified 17 immune subpopulations across lymphoid and myeloid compartments. T cells segregated into CD4+ and CD8+ naïve (CD45RA-CD45RO-) and memory subsets (CD45RA-CD45RO+), as well as a minor double-negative and a phenotypically ambiguous T-cell cluster. B cells resolved into naïve (CD27-) and memory (CD27+) subsets with an additional transitional/atypical cluster expressing high levels of CD20 and CD38. NK cells were separated into CD16+ and CD16− populations, together with a minor heterogeneous NK cluster that expressed CD3. The myeloid compartment comprised classical (CD16-) and non-classical monocytes (CD16+), conventional (CD123-) and plasmacytoid dendritic cells (CD123+), and a small residual myeloid cluster not assignable to canonical subsets.

The cell populations/clusters were visualised using the UMAP algorithm [e7] implemented in R, and heatmaps of median marker expression were generated using the pheatmap R package. Population structure and annotation were summarised in a schematic of the hierarchical gating and clustering strategy (eFigure 4) and further visualised using UMAP overlays of canonical lineage markers (eFigure 5). The number of cells retained per population and time point after QC is provided (eFigure 6) Global variation in immune composition across time points was visualized by PCA using per-sample relative abundances of the 17 populations (eFigure 7).

**Differential abundance analysis of cell populations**

Differential abundance (DA) analysis of annotated mass cytometry populations was performed using the diffcyt framework implemented in R [e8]. The diffcyt-DA-edgeR function was applied using default settings (no trend estimation), with a minimum threshold of ≥ 8 cells per cluster in at least 6 patients to ensure robust statistical testing. Independent DA analyses were conducted for each longitudinal comparison (baseline vs 100 days, 6 months, 12 months, and 24 months), using a paired matrix design including subject ID as a blocking factor to account for within-individual repeated measurements. P-values were adjusted for multiple testing using the Benjamini–Hochberg false discovery rate (FDR) procedure. Statistical significance was defined as FDR-adjusted $p < 0.05$. $Log_2$ fold changes ($log_2FC$) were calculated using the edgeR model and visualized using volcano plots. For visualization purposes, vertical thresholds were drawn at the empirical 10th and 90th percentiles of the pooled $log_2FC$ distribution (−1.25 and +1.97) to highlight the most extreme abundance shifts within the cohort. These thresholds were not used for statistical inference.

**Definition and temporal projection of functional cellular states**

To fully leverage the functional resolution of the mass cytometry dataset, we implemented a workflow combining unsupervised clustering, meta-clustering, and supervised learning to define and track reproducible cellular phenotypes over time. We refer to these phenotypic entities as "cellular states" rather than clusters because they represent stable, multivariate functional phenotypes defined by additional maker expression profiles that can be projected onto newly acquired samples using supervised modelling, rather than re-derived by de novo clustering in each dataset. This allows for new longitudinal samples to be included without running all samples together.

**Discovery cohorts and temporal extension set**

Samples were divided into a discovery cohort for model training and a temporal extension set for model extension. The discovery cohort included samples collected at baseline (BL), 100 days, 6 months, and 12 months. The temporal extension set comprised later time points from the same individuals (24 months) and healthy controls (HCs) as an external reference. All samples underwent identical preprocessing, transformation, and batch control procedures prior to lineage-level clustering and identification of the 17 immune cell populations. The supervised projection step described below was applied only at the level of functional cellular states and did not involve re-clustering of the projection set samples. Importantly, the 24m and HC samples were not used to define cellular states but were used to assess the out-of-distribution stability of the predefined discovery-set states.

**Unsupervised discovery of functional cellular states**

Within the discovery cohort, a random subset of 50,000 cells was sampled across patients and time points. Unsupervised clustering was performed separately within each of the 17 lineage-defined immune cell populations, using FlowSOM followed by meta-clustering with ConsensusClusterPlus (elbow method). Twenty-eight additional surface markers from our antibody panel were used to define cellular states, including adhesion molecules, chemokines receptors, and activation markers (CD235ab, CD133, CD162, CD185, CD278, CD194, CD161, CD184, CD27, CD44, CD127, CD10, CD73, CD146, CD117, CD105, CD49d, CD25, CD49f, CD90, CD195, CD196, CD135, HLA-DR, CD38, CD16, CD33, and CD123). This approach identified 107 distinct cellular states across the 17 immune populations.

**Stability assessment of state definitions**

To quantify the impact of random sampling, the 50,000-cell sampling and clustering workflow was repeated 10 times independently. Reproducibility was assessed by cross-comparing the state centroids (median marker expression profiles) across runs.  High concordance of centroids indicated stable and robust cellular state definitions, minimising sampling-driven stochasticity and ensuring reliability of downstream analyses.

To generalise our findings to unseen data, we trained cell-type-specific LDA models using the MASS R package. First, the 50,000-cell subset was randomly split into two disjoint sets, each containing 25,000 cells. One set was used to train the LDA models, while the other was reserved for testing LDA performance using standard metrics, including sensitivity, specificity, accuracy, and F1 score. A posterior probability threshold of 0.6 was applied throughout to ensure reliable LDA predictions. This internal training-testing procedure within the 50,000-cell subset yielded median sensitivity, specificity, accuracy, and F1 scores of 89.7%, 98.4%, 98.1%, and 88.6%, respectively (eFigure 10a).

**Projection to the remaining discovery data and temporal projection**

Next, LDA models were trained on the full 50,000-cell subset and applied to (i) the remaining cells from the discovery cohort and (ii) the temporal extension cohort (24m and HCs). As ground-truth state labels are not available outside the discovery subset, model consistency was evaluated by computing Pearson correlations between the marker expression profiles of predicted states and the corresponding training-set state centroids. High correlations within the discovery cohort (mean Pearson $r = 0.99$) confirmed robust state assignment (eFigure 10b). Similarly, high correlations in the evaluation set supported generalisation of the learned state definitions to unseen data (eFigure 11).

**Visualisation of cellular states as functional graphs**

Finally, relationships between cellular states were visualised using functional graphs within the Scaffold framework using the grappolo and vite R packages (Pearson correlation as similarity metric), with force-directed layout (ForceAtlas) and visualisation using ggraph [e9]. Similarity between cellular states was represented as functional graphs in which nodes correspond to states and edges connect state pairs with Pearson correlation >0.9 based on their marker expression profiles (eFigure 14). Model-derived log2FC values for all cellular states across longitudinal comparisons are summarised in ordered dotplots (eFigure 12).

**Identification of persistently altered cellular states**

To identify biologically sustained alterations in functional cellular states following aHSCT, we applied an effect-size-based filtering strategy within the discovery cohort (BL–12m). For each cellular state, model-estimated $\log_2$ fold changes ($\log_2$FC) in relative abundance were calculated at each post-treatment time point relative to baseline.

States were considered persistently altered if their $\log_2$FC values exceeded the empirical 10th or 90th percentile thresholds of the pooled $\log_2$FC distribution (−1.25 and +1.97) and remained beyond these thresholds for at least two consecutive post-treatment time points (≥6 months).

These percentile-derived thresholds were used for biological interpretation and visualization only and were not applied for statistical inference. Cellular states meeting persistence criteria in the discovery cohort were subsequently projected to the independent temporal extension cohort using trained LDA models, without recalibration of thresholds.

**Multivariate projection of single-cell profiles using FreeViz**

To assess global functional reprogramming at the single-cell level, we performed supervised multivariate visualisation using the FreeViz algorithm implemented in the Radviz/FreeViz R packages [e10]. FreeViz projects high-dimensional data into a two-dimensional space by optimising the position and weights of dimensional anchors to maximise separation between predefined classes.

Cells from baseline (BL) and 24 months (24m) post-aHSCT were randomly subsampled with equal representation from each class (n = 5000 cells per class, when available). To account for sampling variability and stochastic effects, the subsampling and projection procedure was repeated 1000 times with replacement. The same 28 functional markers used for cellular state definition were included in the FreeViz projection.

For each run, a FreeViz projection was generated, and the optimised marker anchor positions were recorded. The final visualisations represent aggregated density contours across all runs. Marker contribution to class separation was quantified as the Euclidean distance of each marker anchor from the origin in the 2D projection space. Mean distances (± SE) across 1000 runs were calculated and visualised to assess the relative importance of functional markers in distinguishing BL from 24m.

This analysis evaluates global multivariate differences at the single-cell level and does not redefine cellular states.

**Marker perturbation analysis using SARA**

To quantify marker-level perturbations within persistently altered cellular states, we applied the SARA (Significance Analysis of Response to Activation) score [e11]. SARA evaluates differences between full single-cell marker intensity distributions rather than summary statistics, enabling detection of coordinated induction or inhibition across the entire population.

For each cellular state identified as persistently altered in relative abundance, cells from baseline (BL) and 24 months (24m) post-aHSCT were randomly subsampled to ensure equal representation, and SARA scores were computed using 5,000 permutations to assess statistical significance. This permutation-based framework evaluates whether observed distribution shifts exceed those expected by chance, providing robustness against sampling variability, noise, and small responsive subsets.

SARA was applied downstream of cellular state definition and did not influence clustering or LDA modelling. SARA was computed on arcsinh-transformed marker intensities to maintain consistency with clustering inputs. An in-house R implementation was developed based on the original MATLAB algorithm. SARA scores were interpreted as signed measures of marker distribution shifts (positive = relative induction at 24m; negative = relative inhibition at 24m). Because scores are computed independently within each cellular state, their magnitudes are not directly comparable across states.

SARA scores for all functional markers across persistently altered cellular states are presented in eFigure 15.
